# Regional BOLD variability reflects microstructural maturation and neuronal ensheathment in the preterm infant cortex

**DOI:** 10.1101/2025.07.30.667792

**Authors:** Joana Sa de Almeida, Andrew Boehringer, Serafeim Loukas, Elda Fischi-Gomez, Annemijn Van Der Veek, Lara Lordier, Sebastien Courvoisier, François Lazeyras, Dimitri Van De Ville, Gareth Ball, Petra S. Hüppi

## Abstract

BOLD variability reflects meaningful brain activity, yet its structural and biological correlates during early development remain unknown.

We aimed to **i**nvestigate how BOLD variability evolves in very preterm (VPT) infants, its relationship with cortical microstructure and gene expression, and how it differs from full-term (FT) newborns at term-equivalent age (TEA).

Using resting-state fMRI and multi-shell diffusion imaging, acquired in 54 VPT (longitudinally at 33-weeks GA and at TEA) and 24 FT newborns, we evaluated regional differences in cortical BOLD variability and microstructural maturation, and compared to patterns of gene expression in the fetal cortex, using the BrainSpan dataset.

BOLD variability increased in primary sensory-sensorimotor and proto-DMN regions, accompanied by decreased cortical diffusivity. Gene expression analysis revealed concurrent upregulation of genes mediating gliogenesis and neuronal ensheathment. Compared to FT newborns, VPT at TEA showed decreased BOLD variability and increased cortical diffusivity.

BOLD variability reflects cortical microstructure, mediated by upregulation of gliogenesis and neuronal ensheathment. Interruption of these processes by preterm birth identifies putative mechanisms of preterm brain injury.

## 1. Introduction

Neural dynamics can be measured in a non-invasive, though indirect way, using functional MRI (fMRI) through the Blood Oxygen Level Dependent (BOLD) signal, measured with functional MRI. Resting-state fMRI has been a particularly attractive paradigm to obtain richly structured spontaneous activity. So-called “static” analysis techniques obtain measures that summarize the average behavior over a whole run. However, the “BOLD signal variability,” characterized by its variance, captures the magnitude of moment-to-moment regional variations in BOLD signal (Garrett et al., 2010, 2011, 2013). It is thought to indirectly reflect spontaneous brain activity arising from moment-to-moment fluctuations in neuronal activity (Baracchini et al., 2021; Roberts R.P., 2023). These temporal fluctuations in neuronal activity are thought to contribute to synaptic connectivity, consistent with findings from electrophysiological and EEG studies (Faisal et al., 2008; McIntosh et al., 2010; Misic et al., 2011; Uddin, 2020; Vakorin et al., 2011), and thus to underlie brain function.

In adults, BOLD variability varies across the cortex, with greater variability in association and transmodal regions (Baracchini et al., 2021). Greater BOLD variability has been associated with increased task difficulty and improved cognitive performance (Boylan et al., 2021; Garrett et al., 2013; Protzner et al., 2013; Roberts R.P., 2023), and correlates with electrophysiological measures of neural dynamics (Faisal et al., 2008; McIntosh et al., 2010; Misic et al., 2011; Uddin, 2020; Vakorin et al., 2011). BOLD variability increases significantly during childhood (Wang et al., 2021) and is a strong predictor of age across lifespan (Garrett et al., 2010), decreasing with aging (Baracchini et al., 2023; Garrett et al., 2013; Grady & Garrett, 2014; Rieck et al., 2022). So far, no studies have examined changes in BOLD variability in infants prior to term-equivalent age (TEA).

Additionally, few studies have explored the structural and biological underpinnings of BOLD variability, although evidence suggests it correlates with contiguous white matter (WM) microstructural integrity, both in adults and children (Burzynska et al., 2015; Good et al., 2020; Wang et al., 2021). Recently, using the ex-vivo BigBrain histological dataset (Amunts et al., 2013), Baracchini et al. found that BOLD variability was increased in cortical areas with greater laminar differentiation and higher neuronal density, particularly in granular cortical regions where the distinct layer IV supports efficient sensory processing and a broader range of functional responses (Baracchini et al., 2023). However, to date, no studies have examined the relationship between BOLD variability and intra-cortical microstructure, nor its biological correlates, specifically the underlying gene expression patterns, in adults or infants.

Diffusion-weighted imaging (DWI) measures water diffusion in brain tissue at the mesoscopic scale (Basser, 1995), enabling the study of brain microstructure. However, investigating gray matter (GM) microstructure with diffusion MRI (dMRI) poses significant challenges. Unlike white matter (WM), GM does not display orientational coherence, is not as heavily myelinated, and is characterized by a complex intra- and extracellular environment, with multiple barriers and different cellular compartments.

Traditional signal representation models of diffusion, such as Diffusion Tensor Imaging (DTI), provide insights into GM organization by measuring the anisotropic diffusion of water molecules within the tissue, without intrinsic diffusivity assumptions, capturing variations in diffusion patterns (Le Bihan et al., 2001). However, DTI assumes a Gaussian distribution of diffusion, which does not hold true in many biological tissues. To quantify the non-Gaussianity of diffusion, Diffusion Kurtosis Imaging (DKI) has been proposed as an extension of DTI, accounting for the non-Gaussian signal decay and estimating diffusion kurtosis as a reflective marker for tissue heterogeneity (Jensen et al., 2005). Studies have shown that diffusion kurtosis is more sensitive to microstructural complexity within tissues than DTI (Cheung et al., 2009; Paydar et al., 2014). Both DTI and DKI lack specificity for multiple diffusion signals within a single voxel (e.g., crossing fibers) and complex tissue microstructural features. To overcome this, multicompartment models have been proposed, to provide a more complete characterization of tissue complexity. The Spherical Mean Technique (SMT) model estimates microstructural parameters from the direction-averaged diffusion signal, hence removing directionality dependence, and provides measures sensitive to cellular composition, including intra-axonal and extra-axonal compartments (Kaden et al., 2016). Unlike other multi-compartment models, such as NODDI (Neurite Orientation Dispersion and Density Imaging), SMT does not rely on strong assumptions regarding fixed diffusivities, making it more suitable for complex tissue environments, like the GM (Zhang et al., 2012). Taken together, the combination of different dMRI models might provide valuable insights into GM microstructure and its association with BOLD variability.

While imaging can reveal the relationship between BOLD variability and cortical microstructure maturation during early brain development, gene expression analysis can provide mechanistic insight into the processes and pathways underlying developmental functional and structural changes occurring in the cortex. We hypothesized that regional changes in BOLD variability, coupled with changes in the cortical microstructure, are associated with the expression of genes linked to neocortical organization.

In addition to the lack of evidence about how BOLD variability changes during early brain development, few studies have examined the impact of preterm birth on its developmental trajectory (Amunts et al., 2013; Wu et al., 2016). Preterm birth is known to occur during a critical period of activity-dependent plasticity and brain development, characterized by axonal and dendritic growth and arborization, synaptogenesis, neural organization and myelination (Knickmeyer et al., 2008; Kostovic & Jovanov-Milosevic, 2006). Preterm birth exposes the infants to a dramatic environmental change and leads to altered structural and functional brain development with adverse, long-term outcomes (Dubois et al., 2014; Huang et al., 2006; Huppi et al., 1998; Lordier et al., 2019; Nossin-Manor et al., 2013; Sa de Almeida et al., 2023; Sa de Almeida et al., 2020; Volpe, 2001, 2009).

In this study, we aimed to investigate the maturation of BOLD variability in very preterm (VPT) infants, longitudinally, from 33 to 40-weeks gestational age (GA), across different resting-state networks (RSNs). In addition, using a comprehensive dMRI analysis, combining metrics from DTI, DKI, and SMT models, we evaluated how observed changes in BOLD variability across the cortex align with regional measures of cortical microstructure. Furthermore, to better understand the biological correlates of the observed functional and microstructural changes, we examined spatiotemporal patterns of gene expression, in postmortem tissue samples over the same time period. Finally, we aimed to assess weather preterm birth impacts the expected developmental maturation of BOLD variability and cortical microstructure, in comparison to full-term (FT) birth.

## 2. Results

### 2.1. Patient demographics

In both resting-state fMRI (RS-fMRI) and dMRI samples, significant differences between the VPT and FT groups were observed, as expected, in the following perinatal variables: GA at birth, birth weight, birth height, birth head circumference, APGAR at 1 and 5 minutes and incidence of bronchopulmonary dysplasia (BPD). There were no differences between groups in sex, GA at 2^nd^ MRI scan (TEA), neonatal asphyxia, intrauterine growth restriction, intraventricular hemorrhage grade I, and socio-economic parental status score (Largo et al., 1989) (Table 1). There were no significant differences, for any perinatal clinical variable, between the RS-fMRI and dMRI samples, in the VPT infants’ group or the FT newborns’ group.

**Table 1.**
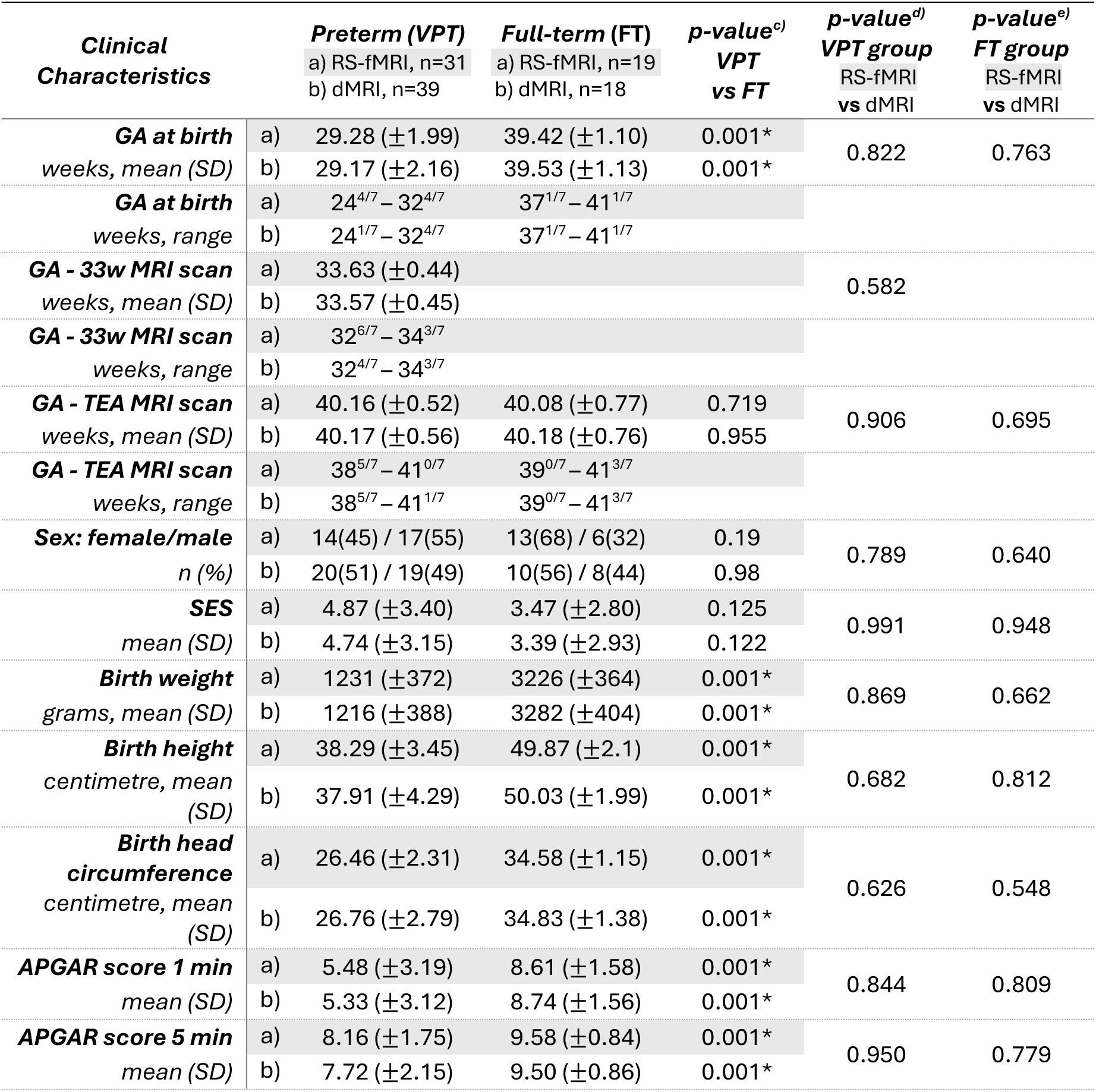
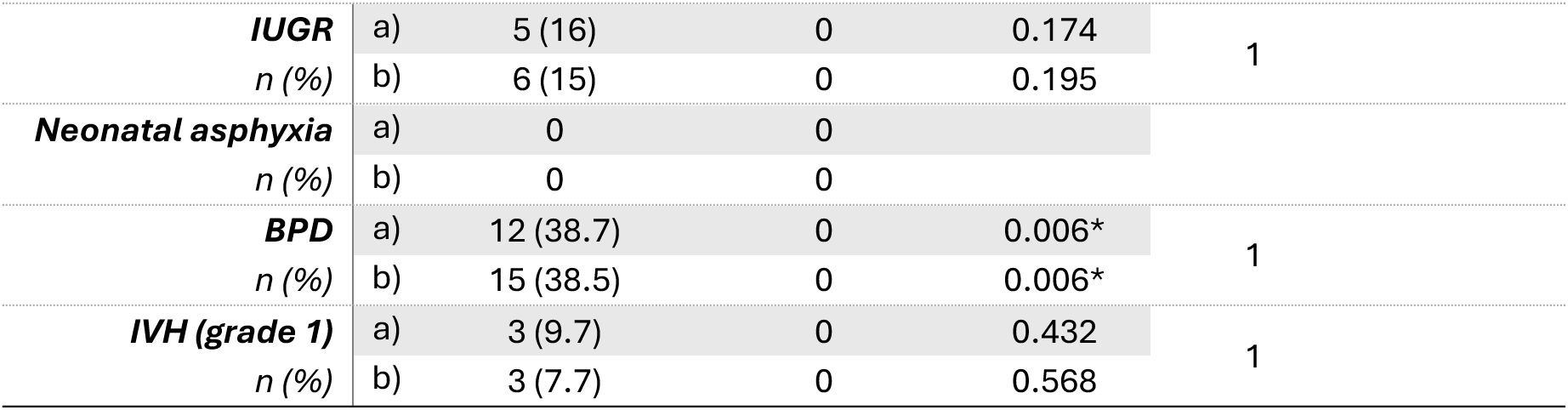
Clinical perinatal characteristics of (a) RS-fMRI and (b) dMRI final samples, for both VPT infants and FT newborns. Group-characteristics per MRI sequence sample [(c) VPT vs FT, within RS-fMRI and dMRI samples], as well as differences between MRI sequence samples [RS-fMRI vs dMRI within each group - (d) VPT and (e) FT], were compared using independent samples t-test for continuous variables and chi-squared test for categorical variables. Significant differences are indicated with (*), p<0.05.

### 2.2. Longitudinal cortical BOLD variability and diffusion microstructural changes during early preterm brain development

In the VPT cohort, BOLD variability increased significantly (false discovery rate (FDR)-corrected; p < 0.05) from 33- to 40-weeks GA (wGA) in the primary sensory (visual, auditory), sensorimotor and proto-Default-Mode-Network (pDMN) (composed by the PCC and precuneus) (cluster 1). BOLD variability did not significantly change in the paralimbic, thalamus, limbic or PFC networks (cluster 2), (Figure 1).

**Figure 1.**
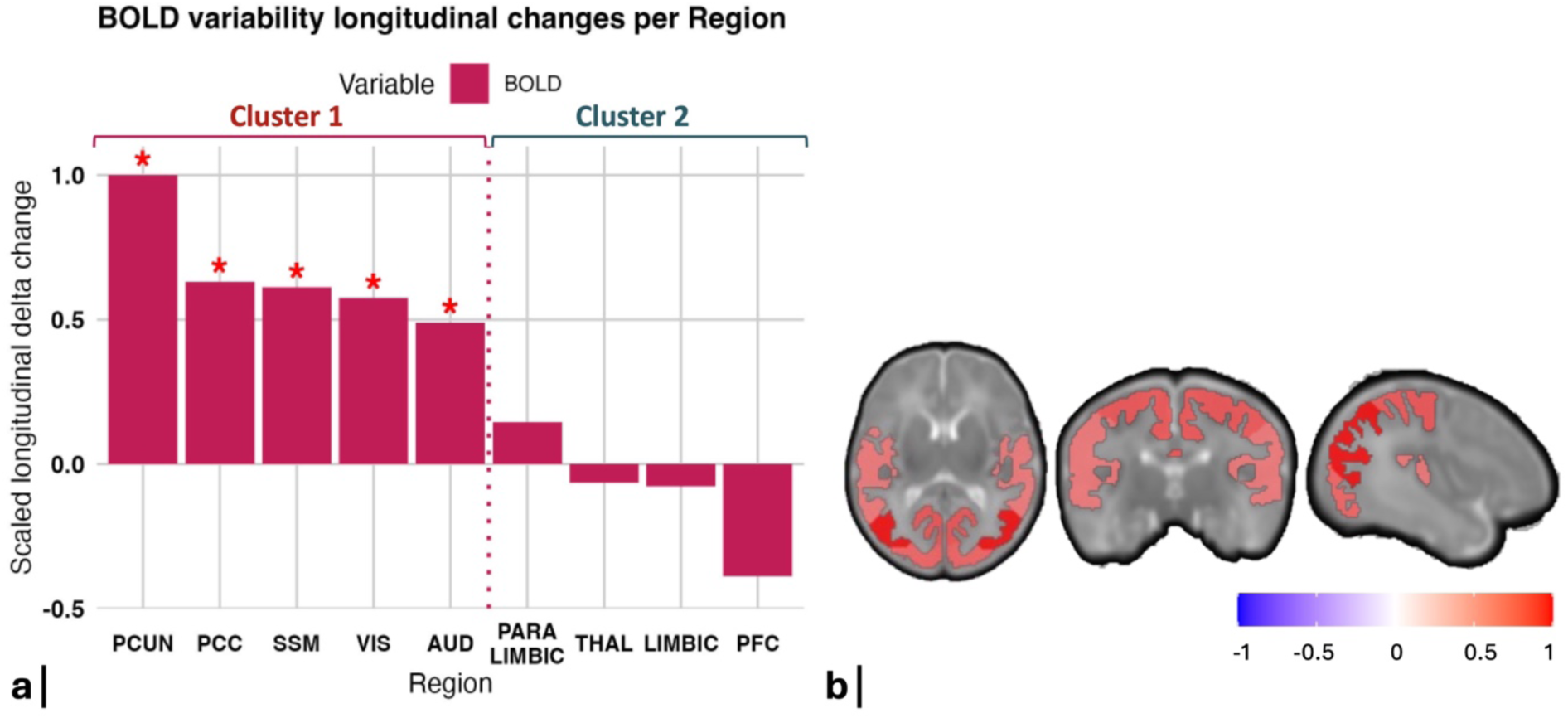
a) Bar plot illustrating BOLD variability longitudinal changes, occurring in VPT infants from 33 wGA to TEA, averaged per RSN and scaled by the largest observed change per metric. Significant changes are highlighted by a “*” (*p*<0.05, after FDR correction). PCUN= precuneus, PCC = posterior cingulate cortex, SSM = sensorimotor, VIS = visual, AUD = auditory, THAL = thalamus, PFC = prefrontal cortex. Regions were clustered according to the presence (cluster 1) or absence (cluster 2) of significant delta changes in BOLD variability. b) Brain plots illustrating the RSNs with significant longitudinal BOLD variability changes occurring in VPT infants from 33 wGA to TEA. The networks include PCUN, PCC, SSM, VIS and AUD, with color mapping representing the magnitude of change.

We next examined changes in diffusion microstructural metrics in VPT infants, from 33- to 40-wGA, across all cortical regions. Significant increases in BOLD variability were associated with an overall decrease in cortical diffusivity and kurtosis (cluster 1, Figure 2).

**Figure 2.**
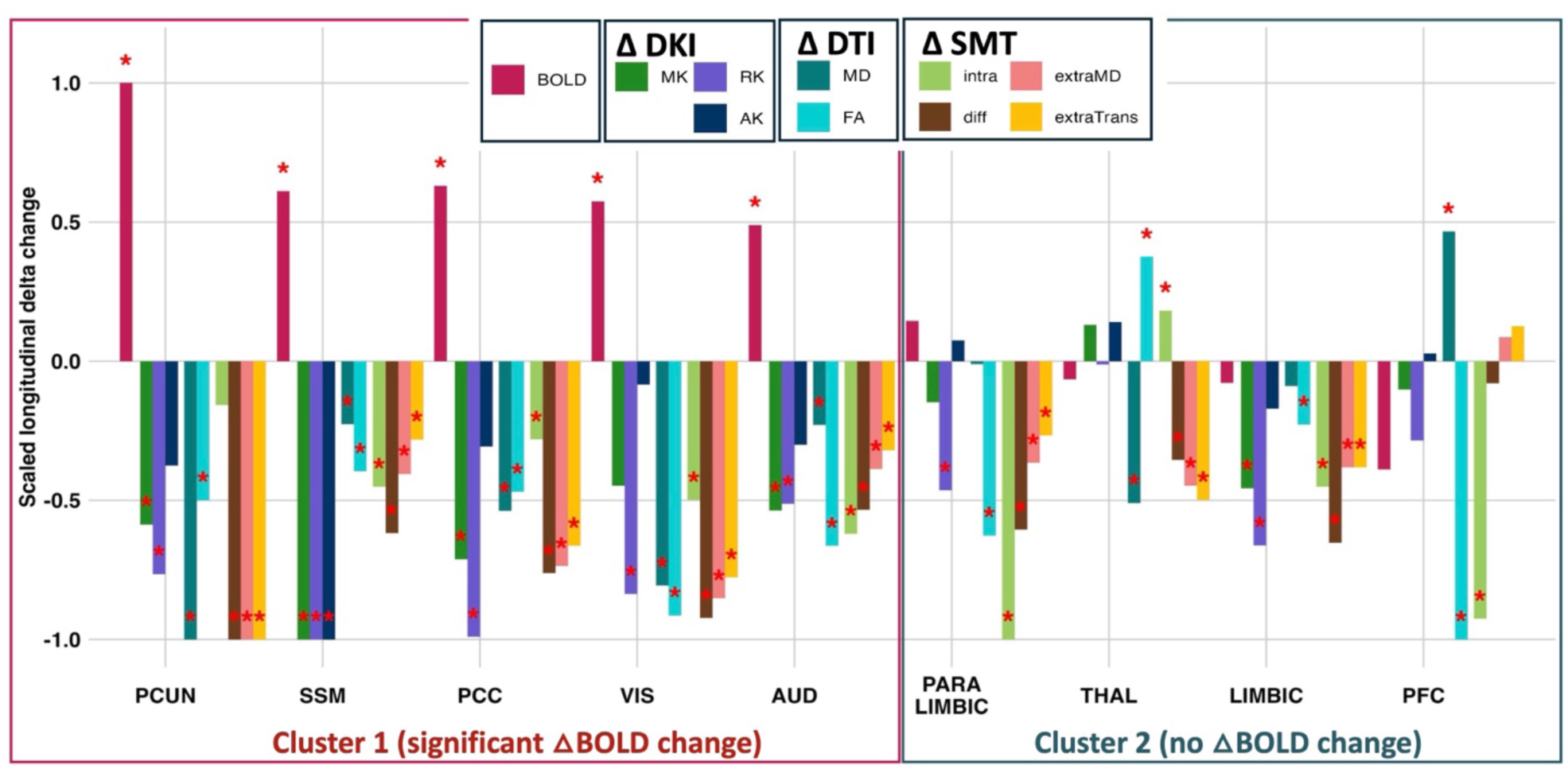
Longitudinal changes in premature infants’ BOLD variability and microstructural diffusivities, from 33- to 40-wGA, averaged per RSN and scaled by the largest observed change per metric. Microstructural DKI metrics include mean (MK), radial (RK) and axial kurtosis (AK); DTI metrics include mean diffusivity (MD) and fractional anisotropy (FA); and SMT metrics include intra-neurite volume fraction (intra), intrinsic diffusivity (diff), extra-neurite mean diffusivity (extraMD) and extra-neurite transverse diffusivity (extraTrans). Significant changes are highlighted by a “*” (p<0.05, after FDR correction). Regions were clustered according to the presence (cluster 1) or absence (cluster 2) of significant delta changes in BOLD variability.

In cluster 1, where BOLD variability increased significantly, SMT metrics globally decreased, namely intrinsic diffusivity (diff), extra-neurite mean diffusivity (extraMD), extra-neurite transverse diffusivity (extraTrans) and intra-neurite volume fraction (intra). Additionally, DTI showed significant decreases of both fractional anisotropy (FA) and mean diffusivity (MD) in all these regions. DKI revealed significant decreases in radial kurtosis (RK) and/or mean kurtosis (MK) across these cluster 1 regions, with axial kurtosis (AK) decreasing specifically in the sensorimotor cortex, in addition to decreases in MK and RK.

In cluster 2, where BOLD variability did not change significantly, the microstructural changes were less coherent. More specifically, the thalamus displayed a significant increase in FA and intra-neurite volume fraction, accompanied by significant decreases in diff, extraMD, extraTrans and MD between 33- and 40-wGA. In contrast, the PFC displayed a marked decrease in both FA and intra-neurite volume fraction, accompanied by an increase in MD. Both limbic and paralimbic regions displayed an overall decrease in diffusivity and/or kurtosis with decreased FA and intra-neurite volume fraction (Figure 2). Additional figures, comprising microstructural metrics longitudinal changes grouped per diffusion technique, can be found in Supplementary Figure S1.

We conducted an additional analysis, clustering cortical regions based on the similarity of their longitudinal microstructural maturation, rather than BOLD variability, to compare the microstructural clusters with those derived from BOLD changes. We found that the microstructural patterns are consistent with the functional partition (Supplementary Figure S2).

#### 2.2.1. Relationship between longitudinal cortical microstructural changes and BOLD variability delta changes

To identify the microstructural metrics that best predicted the observed changes in BOLD variability, we employed a cross-validated LASSO regression model. Out of the 9 microstructural metrics, 3 significant features (MD, RK and MK) were selected as predictors of BOLD variability changes from 33-to 40-wGA. Together, the longitudinal changes in MD, RK and MK accounted for 63% of the observed changes in BOLD variability (R^2^= 0.63). MD showed the strongest association (β = 0.337), followed by MK (β = 0.169) and RK (β = 0.129). An illustration of the selected microstructural delta changes (MD, RK and MK) contributing most significantly to the increases observed in BOLD variability is displayed in Figure 3.

**Figure 3.**
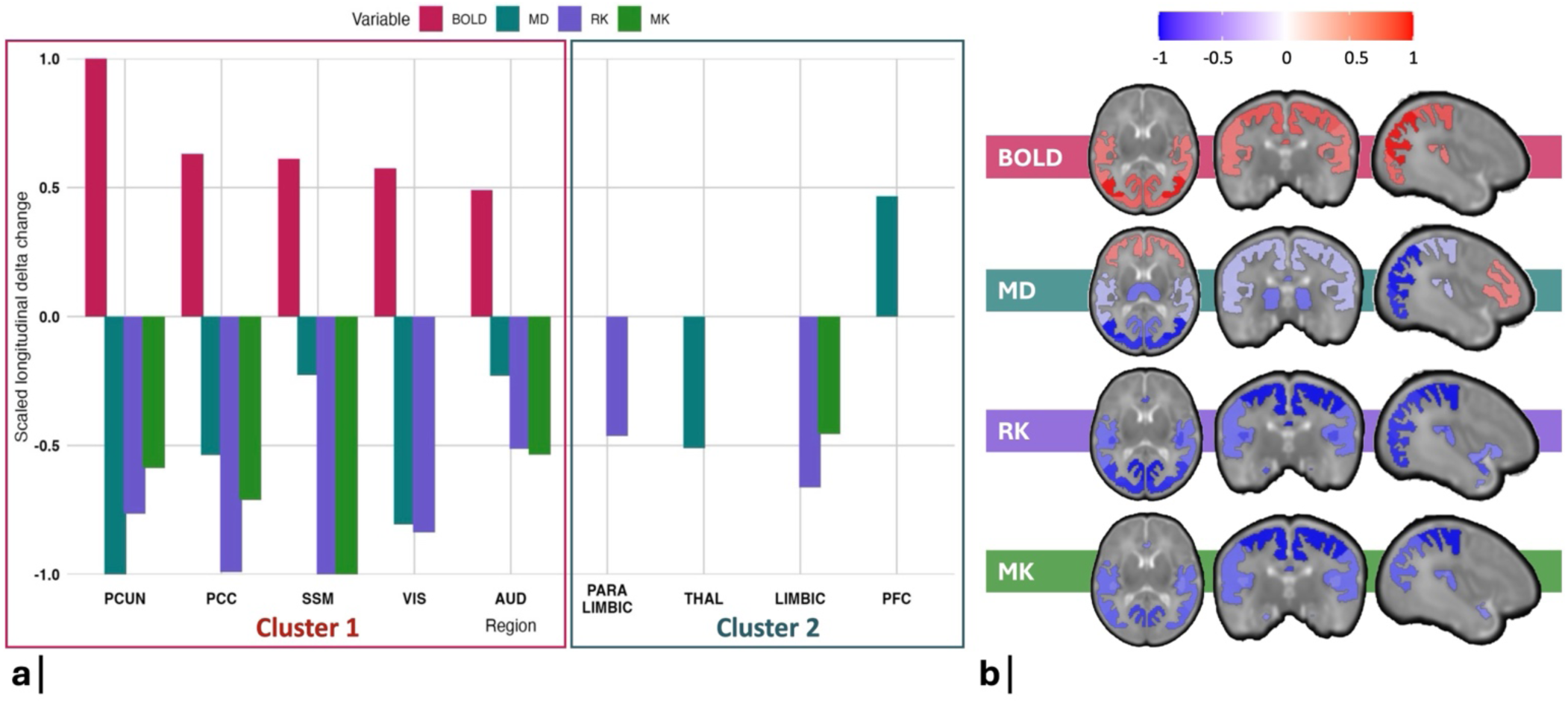
a) Bar graph showing the comparison between the scaled significant longitudinal microstructural changes in MD, RK and MK and the scaled significant longitudinal BOLD variability changes over time. These microstructural features (MD, RK and MK) were identified using Lasso regression as significant predictors of the BOLD variability delta changes. B) Brain plots illustrating the location of the RSNs undergoing significant BOLD variability longitudinal changes, as well as significant changes regarding the MD, RK and MK microstructural metrics. The colormap reflects the magnitude and the direction of the change (increase in red, decrease in blue).

### 2.3. Genetic expression patterns during early brain development

Using a developmental transcriptomic dataset of bulk tissue mRNA data sampled from cortical tissue in 18 post-mortem prenatal human specimens aged 8-37 post conceptional weeks (pcw) (http://development.psychencode.org/), we compared genetic expression patterns between cluster 1 (regions where BOLD variability increased significantly from 33- to 40-wGA) and cluster 2 (regions with no significant longitudinal BOLD variability changes), from mid- to late-fetal period. We hypothesized that regions with changes in BOLD variability (cluster 1), which also presented coupled cortical microstructural maturational changes, would display an increase in expression of genes linked to neocortical organization.

From mid- to late-fetal period (16-22 to 35-37 pcw), 786 out of the total 5,287 genes displayed a significant cluster*time interaction, after correction for multiple comparisons (FDR-corrected; p < 0.05). From these genes, using pairwise comparisons to filter those with differential expression between cluster 1 and 2 from mid- to late-fetal, we identified 132 genes that increased significantly more in cluster 1 vs 2, whereas only 9 genes increased significantly more in cluster 2 vs cluster 1. Additionally, 64 genes were found to decrease significantly more in cluster 1 vs 2, and 45 genes decreased significantly more in cluster 2 vs 1.

Using geneset enrichment analysis (Wang et al., 2017), with a background set of 5,287 genes (Ball et al., 2020), we identified significant enrichments for several important neocortical organizational processes in genes that increased significantly more in cluster 1 than 2. These included: “neuronal ensheathment” (FDR = 0.000014, enrichment = 6.15, 14 genes), “gliogenesis” (FDR = 0.018, enrichment = 2.85, 14 genes, from which 7 genes are common to neuronal ensheathment), “extracellular structure organization/external encapsulating structure organization” (FDR = 0.018, enrichment = 3.58, 10 genes), “cytokine-mediated signaling pathway” (FDR = 0.008, enrichment = 3.6, 13 genes), “coagulation/wound healing” (FDR = 0.008/0.01, enrichment = 4.09/3.03, 10/14 genes, where the 10 genes from coagulation are common to both categories), “extrinsic apoptotic signaling pathway” (FDR = 0.01, enrichment = 3.8, 10 genes) and “muscle cell proliferation (FDR = 0.02, enrichment = 3.8, 9 genes) (Figure 4c). A list of all significant categories and their assigned genes can be found in Supplementary Table S3. No significant enrichment analysis results were found for the genes that increased significantly more in cluster 2 than 1, or for the genes that decreased significantly differently between cluster 1 and 2.

**Figure 4.**
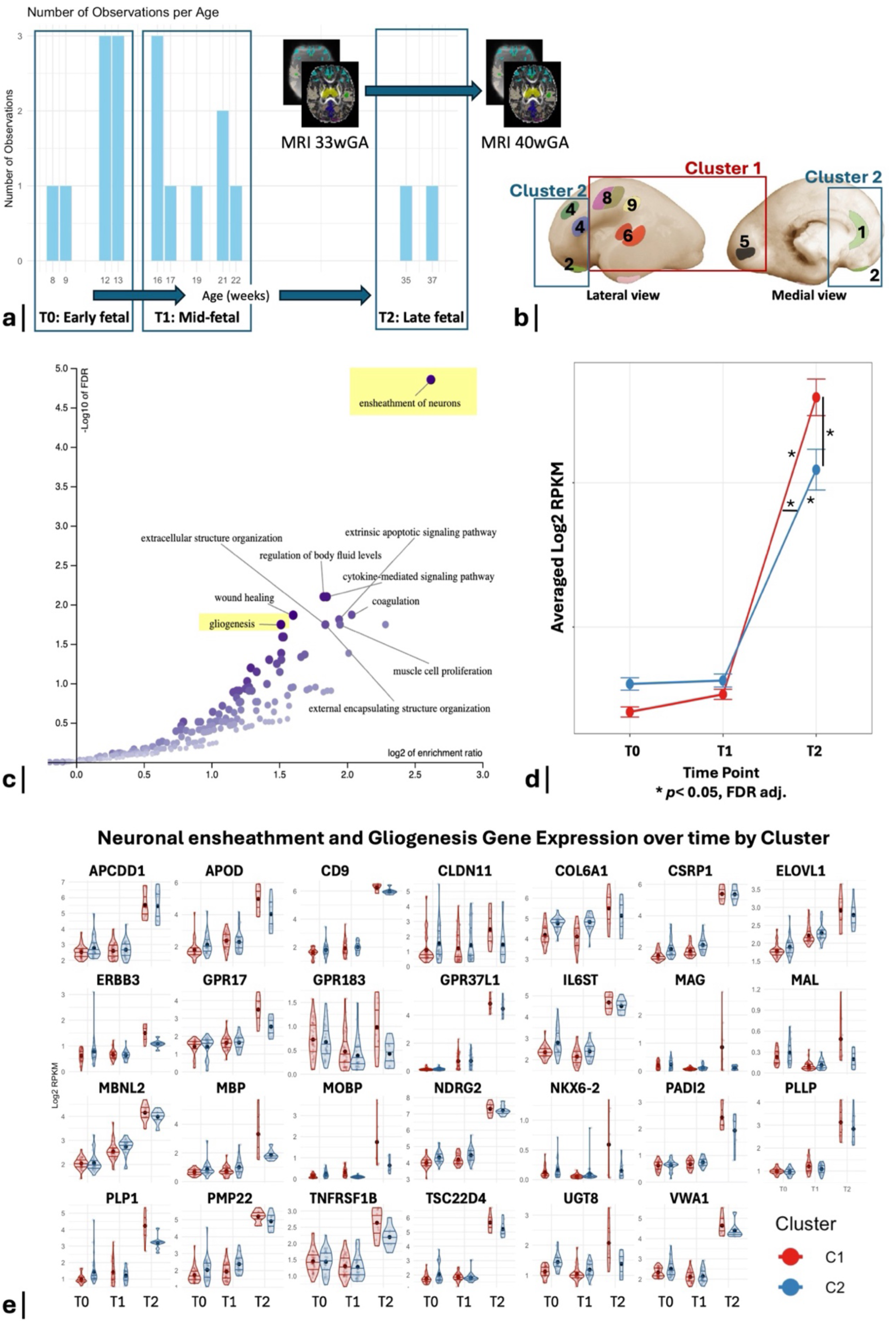
a) Bar plot showing the number of observations (specimen ID) per gestational age (in weeks), obtained from the BrainSpan developmental transcriptomic dataset (http://development.psychencode.org/). The transcriptomic data was clustered per period of gestation (T0: early-fetal, 8-13 pcw; T1: mid-fetal, 16-22 pcw; T2: late-fetal, 35-37 pcw). The longitudinal brain MRI scans were acquired from our preterm infants’ cohort at a first time-point between the mid- and late-fetal periods (33^th^ wGA) and a second one just after late-fetal period (40^th^ wGA). b) Topographic locations of the samples from the BrainSpan developmental transcriptomic dataset. Schematic shows anatomical positions of dissections of the prenatal brain samples, which are coded according to their correspondence to our RSNs brain atlas: (1) medial frontal cortex (MFC) = Limbic, (2) orbitofrontal cortex (OFC) = Paralimbic belt, (4) dorsolateral and ventrolateral prefrontal cortex (DLPFC+VLPFC) = PFC, (5) primary visual cortex (V1) = Visual, (6) primary auditory cortex (A1C) = Auditory, (8) primary motor and primary somatosensory cortex (M1+S1) = Sensorimotor, (9) inferior parietal cortex (IPC) = Precuneus. These regions were clustered according to the presence (cluster 1) or absence (cluster 2) of significant BOLD variability changes from 33- to 40-wGA. Image adapted from (Pletikos et al., 2014). c) Volcano plot showing significant enrichment of Gene Ontology terms (biological processes) (FDR-corrected, p-value < 0.05) of the genes expressed significantly higher in cluster 1 than cluster 2 from mid- to late fetal period. d) Line graph representing the combined expression (averaged Log2 RPKM) of the 27 genes belonging to neuronal ensheathment and/or gliogenesis categories per time-point (T0: early-fetal; T1: mid-fetal; T2: late-fetal) and per cluster (1 in red; and 2 in blue). ANOVA with post-hoc Tukey’s test for pairwise comparison was performed. Mean and 95% confidence intervals are illustrated in red for cluster 1 and in blue for cluster 2. Significant results are highlighted by a “*” (FDR-corrected, p<0.05). e) Violin plots illustrating the expression (in Log2 RPKM) of each of the 27 genes belonging to neuronal ensheathment and/or gliogenesis categories across time points (T0, T1 and T2), per cluster (C1 – cluster 1; C2 – cluster 2).

The combined list of enriched genes associated with neuronal ensheathment and gliogenesis categories comprised a total of 27 genes, including constituents of the myelin sheath (*MBP, MAG, MAL, MOBP, PLLP, PLP1, PMP22, CLDN11*), genes with a role in myelination regulation (*NKX6-2, UGT8, GPR17*), important for oligodendrocyte function and expression (*MBNL2, APOD, VWA1*), for tissue integrity (*COL6A1*), with a role in growth factor signaling (ERBB3), neuroprotection and/or glioprotection (*TNFRSF1B, GPR37L1, IL6ST*), neuronal and glial differentiation and development (*GPR183, APCDD1, NDRG2, CSRP1, TSC22D4, CD9*) and in fatty acid and sphingolipid biosynthesis (*ELOVL1, PADI2*). ANOVA followed by Tukey’s test identified a significant increase in expression of this group of genes in both clusters (1 and 2) specifically from mid- to late-fetal (T1 to T2), but not from early-to mid-fetal (T0 to T1). The increase from T1 to T2 was significantly higher in cluster 1, compared to cluster 2. At the late-fetal time-point (T2), these same genes were also expressed at significantly higher levels in cluster 1 compared to cluster 2. In contrast, at T0 and at T1, there were no differences in the expression of these genes between clusters (Figure 4d). The expression of each of these genes at each time-point and per cluster is shown in Figure 4e.

### 2.4. Effect of preterm birth on BOLD variability and cortical microstructure at TEA

Compared to FT newborns, VPT at TEA showed decreased BOLD variability in the thalamus, PCC, visual and auditory cortices. Regarding cortical microstructural differences, VPT at TEA displayed overall increased diffusivities (MD, extra-neurite mean diffusivity, extra-neurite transverse diffusivity and intrinsic diffusivity) in all RSNs cortical regions, in comparison to FT. VPT-TE also presented increased kurtosis (MK and RK) across all RSNs cortical regions, apart from the thalamus (Figure 5). There were no significant differences between groups regarding AK across all cortical regions. Details of all regional differences per diffusion metric, between VPT and FT infants, are provided in Supplementary Table S4.

**Figure 5.**
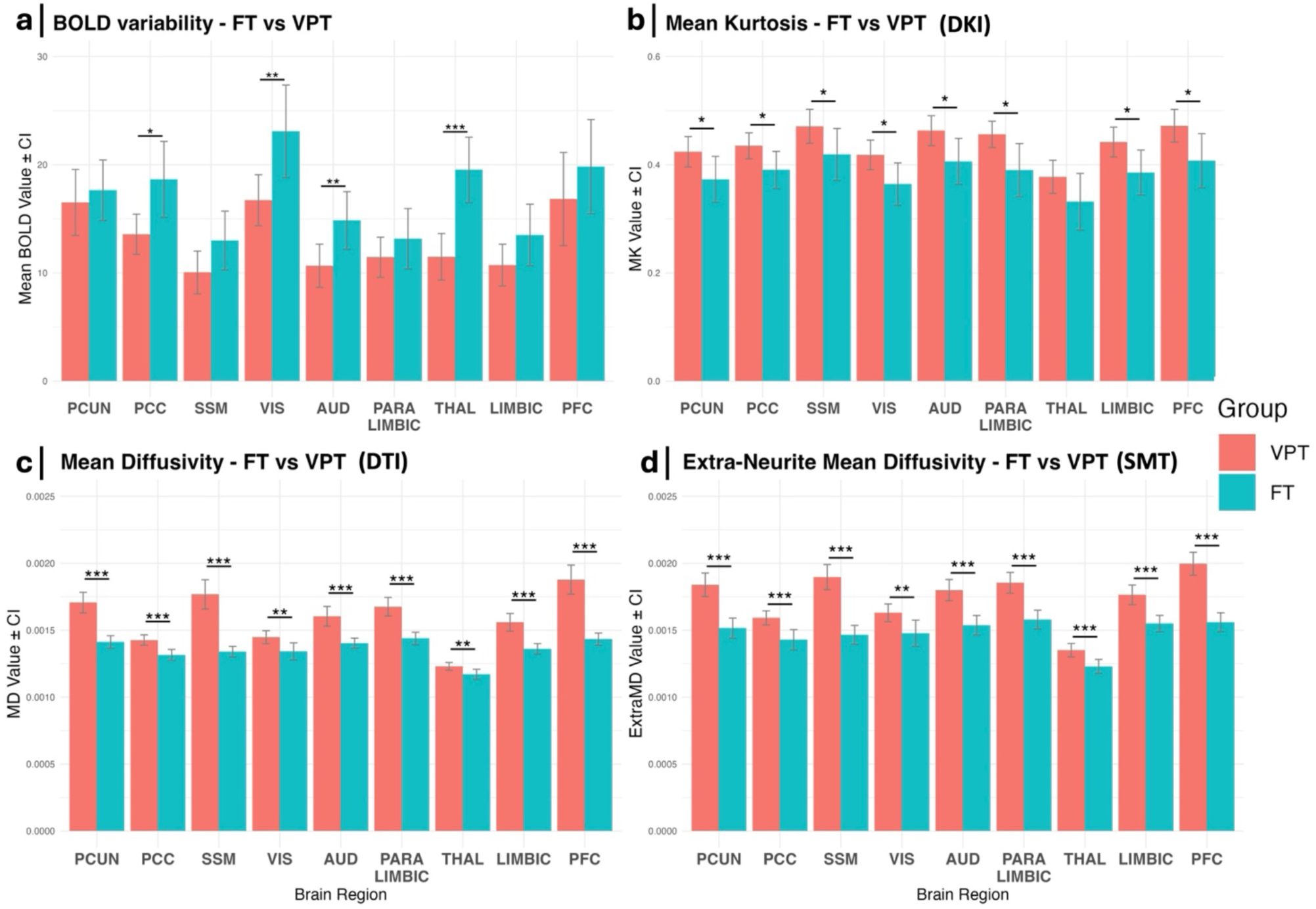
Effect of preterm birth on BOLD variability and cortical microstructure at TEA. Bar plots illustrating mean (with 95% confidence intervals) of BOLD variability (a), Mean Kurtosis (b), Mean Diffusivity (c) and Extra-Neurite Mean Diffusivity (d) in FT infants (blue) and VPT infants at TEA (red) per RSN. Lines indicate significant differences between groups with *p < 0.05, **p < 0.01, ***p < 0.001, FDR adjusted. PCUN = precuneus, PCC = posterior cingulate cortex, SSM = sensorimotor, VIS = visual, AUD = auditory, THAL = thalamus, PFC = prefrontal cortex.

## 3. Discussion

In this study, we combined in vivo fMRI and dMRI data with independent ex vivo gene expression analyses to explore the regional maturation trajectory of BOLD variability across RSNs in very preterm infants, longitudinally, from 33wGA to TEA. Our findings reveal an alignment between brain developmental changes in BOLD variability and regional cortical microstructural maturation, as assessed through advanced dMRI models, including DTI, DKI, and SMT. Furthermore, we identified specific spatiotemporal patterns of gene expression that may underlie the observed changes.

Additionally, using a cross-sectional approach at TEA, we demonstrate the impact of preterm birth on the typical developmental trajectory of BOLD variability and cortical microstructure, in comparison to full-term birth.

### 3.1. Cortical microstructural maturation underlies RSNs BOLD variability increases during preterm infants’ brain development

From 33wGA to TEA, BOLD variability significantly increased in preterm infants, specifically in the primary sensory (visual and auditory), sensorimotor and proto-DMN (precuneus and PCC) (cluster 1). The primary visual, auditory and somatosensory regions are known to present an early establishment of activity-dependent thalamo-cortical connectivity. In fact, from 24-to 32-post conceptional weeks, thalamocortical fibers grow into and accumulate in the subplate, beginning in somatosensory areas (Kostovic & Rakic, 1990), followed by the visual (Kostovic & Rakic, 1984) and auditory (Krmpotic-Nemanic et al., 1983) cortices, and leading to the formation of synapses in the deep cortical plate during prenatal development (Molliver et al., 1973). During this period, electrophysiological phenomena can be detected by EEG, with the appearance of evoked potentials and giant transients, indicating synaptogenesis, cortical activity and processing of sensory information (Hrbek et al., 1973; Vanhatalo et al., 2005). Indeed, during brain development, synaptogenesis was shown to follow a spatiotemporal pattern, occurring earlier in primary motor and sensory areas, and later in the prefrontal cortex (Huttenlocher & Dabholkar, 1997). Our findings, revealing significant longitudinal increases in BOLD variability specifically in primary motor, sensory, and posterior brain regions from 33wGA to TEA, with no changes in prefrontal regions, agree with the spatiotemporal pattern of synaptogenesis and cortical activity.

Regional functional BOLD variability increases in cluster 1 were accompanied by significant decreases in cortical microstructural diffusivity and kurtosis, as well as by decreases in FA and intra-neurite volume fraction, evaluated using different dMRI models (DTI, DKI and SMT).

Decreases in cortical diffusivity (mean diffusivity, intrinsic diffusivity, extra-neurite mean diffusivity) from 33wGA to TEA, align with an ongoing microstructural cortical maturation, possibly relating to an increased dendritic arborization, formation of basal dendrites cross-connections and gliogenesis, that occur during this period (Bystron et al., 2008; Cameron & Rakic, 1991; Marin-Padilla, 1992; Rakic, 2003). Previous studies support the notion that MD decreases in the human cortex throughout preterm development (Ball, Srinivasan, et al., 2013). Decreases in cortical MD have been linked to an increase in dendritic density and synaptic complexity, as observed in histological studies of rodents (Sizonenko et al., 2007). In addition, we have previously shown that these regions in cluster 1, namely the visual, auditory, sensorimotor, precuneus and cingulate cortices, undergo a significant increase in diffusion ODI (orientation dispersion index) and FC (fiber cross-section) from 33wGA to TEA, further supporting an increase in cortical complexity and cellular density during this period (Sa de Almeida et al., 2023).

The observed decrease in cortical mean kurtosis is consistent with previous findings evaluating human brain prenatal development (Ouyang et al., 2019). This is in contrast with the known increase in mean kurtosis in the cortex observed later, during postnatal period and childhood, especially during the first 2 years of life (Paydar et al., 2014; Shi et al., 2019; Wang et al., 2023). The observed decrease in mean kurtosis during the prenatal period was hypothesized to indicate a continuous decrease in diffusion barriers, possibly caused by a decrease in neuronal/cell density during this period (Leuba & Garey, 1987; Steven et al., 2014; Wu & Cheung, 2010). In agreement with this theory is our observation of a longitudinal decrease of intra-neurite volume fraction across these cortical regions, which is in line with previous studies showing a decrease in neurite density index (NDI) across grey matter tissue during this period (Batalle et al., 2019). Decreases in cortical neuronal density could be explained by synaptic pruning and physiological apoptosis, involution of the subplate, as well as by a cortical expansion, which may lead to an apparent decrease in neuronal density (Rabinowicz et al., 1996). Additionally, decreases in intra-neurite volume fraction may be led by the disruption of the radial glial scaffolding, known to occur during this period of development, and contribute to a reduction of cortical cell density (Bystron et al., 2008; McKinstry et al., 2002). It is unlikely that measures such as intra-neurite volume fraction and NDI solely reflect neurite structures, with possible contributions from prominent radial glial fibers, which exhibit elongated, coherent radial organization, and may be interpreted as intra-neurite compartments.

Interestingly, we show that the decreases in MK are mostly led by decreases in RK, and not AK. These kurtosis reductions, particularly in the radial direction, support that disruption of the radial glia scaffolding may be also at the origin of the observed mean kurtosis changes. Notably, the sensorimotor cortex was the only region where all kurtosis metrics (MK, RK and AK) decreased significantly and in a more pronounced manner than all the other diffusivity measures. The AK decrease in this region suggests that diffusion barriers decrease not only in the radial direction, as in other cortical areas, but also in the direction of the axonal fibers. The sensorimotor cortex has been shown to be the first cortical region to present myelin, among all the others, during brain development (Miller et al., 2012). Although myelin has not been histologically demonstrated to be present in the human cortex at birth, myelination of white matter fibers starts during the second trimester of pregnancy in the corticospinal tract and is known to increase in response to electrical excitation (Wake et al., 2011). The sensory input to the sensorimotor cortex may therefore contribute to the myelination of the corticospinal tract and lead to pre-myelination events in the sensorimotor cortex, possibly resulting in early axonal ensheathment by pre-myelinating oligodendrocytes. In fact, the initiation of intracortical myelination is thought to involve the maturation of oligodendrocytes, which begin to form myelin sheaths around axons within the deeper layers of the cortex, and to start in sensorimotor regions compared to association areas (Call & Bergles, 2021).

The overall decreases in cortical diffusivity and in intra-neurite volume fraction were accompanied by decreases in FA in cluster 1 regions. The FA reduction is in alignment with previous studies showing decreased cortical anisotropy during prenatal development, attributed to reduced radial glia scaffolding and increasing dendritic complexity (Ball, Srinivasan, et al., 2013; Huang et al., 2013; McKinstry et al., 2002; Neil et al., 1998; Sizonenko et al., 2007).

Cluster 2 cortical regions, where BOLD variability did not change significantly from 33-wGA to TEA, inhibited heterogenous diffusion changes, with less pronounced diffusivity decreases. In particular, the PFC and the paralimbic region (combining the OFC and the temporal pole) showed marked FA and intra-neurite volume fraction reductions, exceeding diffusivity changes. Literature indicates that frontal association areas have lower dendritic density, synapse number and glia density than primary sensory regions in neonates (Huttenlocher & Dabholkar, 1997; Jakovcevski & Zecevic, 2005; Travis et al., 2005). This could explain the less pronounced decreases in diffusivity in frontal areas. Furthermore, the marked decline in FA and NDI in frontal regions has been consistently observed in other human prenatal studies, and has been shown to correlate with increased grey matter volume and mean curvature (Ball, Srinivasan, et al., 2013; Batalle et al., 2019; Yu et al., 2016). This suggests that cortical volume expansion would be driven by dendritic arborization and neuropil expansion, rather than the addition of new cells (Bayer & Altman, 1991). When volume expansion outpaces the cellular density increases, FA and intra-neurite volume fraction decline, while MD may increase, as observed in the PFC. These microstructural changes indicate overall delayed maturation in more frontal regions, potentially explaining the absence of significant BOLD variability increases in these regions.

Interestingly, the thalamus showed increased FA and intra-neurite volume fraction with decreased diffusivity. In fact, as a subcortical region, the thalamus exhibits a distinct microstructural organization in comparison to cortical grey matter areas, which can explain these findings. Despite the important establishment of thalamocortical connectivity during this period (Katz & Shatz, 1996; Kostovic & Judas, 2010; Mizuno et al., 2010), no significant increase in BOLD variability was observed in this region.

To better understand which diffusion metrics most effectively explain the observed regional increases in BOLD variability, we performed a Lasso regression analysis. This analysis identified longitudinal decreases in MD, MK and RK as the key predictors, together accounting for up to 63% of the observed changes in BOLD variability. This provides evidence for the importance of cortical microstructure maturation in enabling functional activity. Decreases in these diffusivity metrics, and in particular the strong association with the decrease in MD, support that, possibly, an increase in dendritic arborization, dendritic density, synaptogenesis and gliogenesis (Sizonenko et al., 2007) may contribute to BOLD variability increase. In fact, the intracortical burst of dendritic growth and maturation is known to occur in parallel with rapid synaptogenesis during the late mid-fetal and early late-fetal periods, coinciding with changes in functional properties of the neuronal cortical circuitry and appearance of evoked potentials (Brodmann, 1909; Silbereis et al., 2016).

Furthermore, the association of increased BOLD variability with decreases in MK and RK denote a decrease in diffusion barriers, possibly due to the disruption of the radial scaffold with the ongoing cortical maturation, allowing the extension of dendritic arborization, and also its differentiation into more localized glia cells once cortical migration finishes, ultimately supporting synaptogenesis (Rakic, 1988; Volpe, 2019). In fact, literature suggests that regions where radial glia disappear earlier tend to exhibit earlier synaptic activity (Molnar et al., 2019; Noctor et al., 2001; Rakic, 1988).

### 3.2. Genes mediating gliogenesis and neuronal ensheathment increase more in RSNs where BOLD variability increased

Between 16-22 pcw and 35-37 pcw, corresponding to the mid-to late-fetal period, analysis of ex-vivo fetal cortical gene expression patterns revealed a significant upregulation of genes associated with neuronal ensheathment and gliogenesis. This increase is significantly higher in the regions where BOLD variability increases from 33-to 40-wGA, namely the primary sensory (visual and auditory), sensorimotor and proto-DMN (precuneus and PCC). The expression of these genes remains significantly higher in these same regions during the late-fetal period, compared to regions without significant BOLD variability changes.

This prenatal regional upregulation of genes involved in neuronal ensheathment and gliogenesis coincides temporally and spatially with the functional and microstructural cortical maturation observed via MRI, characterized by longitudinal increases in BOLD variability, and reductions in cortical diffusivity and kurtosis.

A previous study in an independent dataset demonstrated a spatial alignment regarding cortical maturational differences between primary and higher-order cortical regions in term born infants, using MRI-based measures (T1/T2 ratio, cortical thickness, FA, MD, ODI and NODDI intra-cellular volume fraction), reporting that the spatial patterning of cortical gene expression during late gestation was mirrored by a regional variation of cortical microstructure at term. More specifically, in comparison to higher-order areas, genes expressed in primary sensory areas at late gestation were found to belong to specific glial populations, including oligodendrocytes and microglia (Ball et al., 2020). This aligns with our findings, denoting an increased expression of genes involved in neuronal ensheathment and gliogenesis in broadly the same regions, further elucidating mechanisms underlying the spatial variation in brain cortical maturation captured by our functional and microstructural MRI data.

Despite the absence of histologic evidence for intracortical myelination in the human fetus and at the time of birth (Kinney et al., 1988), it is possible that an increase in oligodendrocyte precursor cells (OPC) and early axonal cortical ensheathment by immature oligodendrocytes (O4+, O1+) may be at the origin of the observed decreases in cortical microstructural diffusivity. In fact, literature shows that OPC and immature oligodendrocytes produce mRNA levels of genes associated with myelination (such as MBP, PLP1 and MAG), as these cells prepare for differentiation, even before the myelin is detectable by immunostaining (Marques et al., 2016; Zhang et al., 2014). The decreases in kurtosis occurring during the same developmental period, especially in the radial direction, may reflect the retraction and disappearance of radial glia, an event that is known to coincide with the expansion of OPC during late gestation, marking a transition from neurogenesis to gliogenesis (Aquilino et al., 2025; Kostovic et al., 2021; Rowitch & Kriegstein, 2010).

Furthermore, literature supports that the OPC and the early cortical axonal ensheathment by immature oligodendrocytes may both contribute to and be promoted by increased synaptogenesis (Bergles et al., 2010; Fields, 2015; Sakry et al., 2014). This interplay could explain the concomitant increase observed in BOLD variability within the same cortical regions where the expression of these cells is increasing.

This analysis therefore provides important insights regarding the cellular processes underlying both the diffusion microstructural changes as well as functional changes observed with in vivo MRI data during this key period of brain development.

### 3.3. Preterm birth impacts BOLD variability and cortical microstructure at TEA

Compared to FT newborns, premature infants at term-equivalent age (TEA) showed reduced BOLD-variability specifically in sensory RSNs (auditory and visual), PCC and thalamus, as well as overall decreased microstructural diffusivity and kurtosis across all cortical regions, denoting the impact of preterm birth on cortical functional and structural RSNs maturation.

We found that the cortical microstructure of VPT infants at TEA was characterized by overall increased diffusivity and kurtosis across all RSNs cortical regions. Only a few studies have compared cortical microstructure between full-term newborns and very preterm infants at TEA using dMRI. Our findings align with the works of Ball et al., which demonstrated that, in comparison to FT, preterm infants at TEA present higher MD in frontal, parietal, occipital and temporal cortical regions (Ball et al., 2020; Ball, Srinivasan, et al., 2013). We provide the first evidence of cortical kurtosis differences between these two groups at term-equivalent age, offering new insight into early microstructural development. VPT at TEA presented higher MK across all cortical RSNs regions, except for the thalamus, where there were no differences between groups. Interestingly, these increases in MK were led by increases in RK (with no changes observed in AK between groups). Overall, our results suggest a delayed microstructural cortical maturation in VPT at TEA compared to FT.

As our study and others have shown, cortical diffusivity and kurtosis tend to decrease during brain development, until TEA (Ball, Srinivasan, et al., 2013; Ouyang et al., 2019). As stated before, the longitudinal decreases in cortical diffusivity is thought to relate to an increase in dendritic arborization and density, as well as in gliogenesis, whereas the longitudinal decreases in MK and RK may derive from the disruption of the radial glial scaffolding. These processes thus reflect ongoing cortical maturation, and appear to be disrupted by preterm birth, leading to increased cortical diffusivity and kurtosis at TEA in VPT compared to FT. In fact, cortical neurogenesis is mostly finished by mid-gestation, before 27 wGA, and it is followed by a shift to intense gliogenic events (Kostovic et al., 2021; Silbereis et al., 2016). Preterm birth is known to expose the brain to noxious environmental exposures, conferring vulnerability during these important developmental processes. Prematurity may thus lead to a disruption of the ongoing gliogenic events, particularly oligodendrocyte maturation, which occurs during late gestation. In fact, hypoxic-ischemic events (a model of preterm injury) have been shown to damage precursors of oligodendrocytes (Back, 2006; Tsai et al., 2016). In agreement with our findings and hypothesis, Ball et al., have shown that, compared to FT, preterm infants at TEA present, on average, lower cortical T1w/T2w contrast, and this contrast was found to correlate positively with the expression of genes associated with glial cells, including oligodendrocytes, during the second half of gestation (Ball et al., 2020).

Interestingly, the thalamus was the only region where no significant differences between groups were found for RK and MK. This may be due to the thalamus’s distinct microstructural organization compared to the cortex, being more compact and exhibiting radial glia fibers mainly in the ventral nuclei only, which may reduce the sensitivity of kurtosis to detect group differences (Forutan et al., 2001; Kim et al., 2023). Nevertheless, group differences in diffusivity remained evident, with VPT at TEA presenting higher diffusivity in the thalamus than FT.

Despite the need of further research to better elucidate the mechanisms behind the increased cortical diffusivity and kurtosis observed in VPT at TEA in comparison to FT, our findings combining longitudinal dMRI data and genetic expression analysis during late gestation support that preterm birth may disrupt ongoing gliogenic and pre-myelination events, leading to an altered cortical microstructure by TEA.

This study also provides the first evidence of the impact of preterm birth on BOLD variability, which is decreased in VPT at TEA, namely in sensory RSNs (auditory and visual), PCC and thalamus.

As stated, the rapid synaptogenesis during the late mid-fetal and early late fetal period is known to be heterochronous, following a spatiotemporal pattern, with earlier occurrence in primary motor and sensory areas, and latest in the prefrontal cortex (Huttenlocher & Dabholkar, 1997). It is possible that preterm birth, by disrupting the ongoing maturational processes, namely dendritic arborization and gliogenesis, will consequently impact synaptogenesis in the regions undergoing the most important maturational changes during late gestation, specifically primary sensory areas. Besides the visual and auditory, according to our findings, the PCC was also among the cortical regions undergoing significant longitudinal increases in BOLD variability from 33wGA to TEA, which may make it more vulnerable to preterm birth related injury, in comparison to other cortical regions.

Previous studies evaluating the impact of preterm birth at TEA on functional connectivity have shown that VPT at TEA present a decreased RS-functional connectivity between the salience network and the auditory, visual, PCC and thalamic networks, compared to FT (Lordier et al., 2019). Other studies using fMRI and dMRI have also shown that the visual, PCC and thalamus are regions that are highly connected to each other and to others (rich-club nodes), and present decreased functional and structural connectivity in preterm infants at TEA compared to FT (Sa de Almeida et al., 2020; Scheinost et al., 2016).

In particular, a decreased thalamo-cortical structural and functional connectivity has been consistently shown across studies in preterm infants at TEA compared to FT newborns (Ball, Boardman, et al., 2013; Sa de Almeida et al., 2020; Smyser et al., 2010). In fact, thalamocortical connections are being established during the late second and third trimester of brain development, which is when preterm birth occurs (Kostovic & Jovanov-Milosevic, 2006; Kostovic & Judas, 2010), exposing the brain to noxious events such as inflammation, hypoxia/ischemia and stress, which may affect the establishment of these connections. Disruption of thalamo-cortical connectivity may interfere with synaptogenesis in cortical primary sensory regions (such as the visual and auditory cortex), which rely heavily on the thalamic input for development, and are among the first to receive it. The PCC also receives thalamic input, although this occurs relatively later during gestation compared to primary sensory areas (Kostovic, 1990), which makes it a vulnerable target to the effects of preterm birth.

By linking our imaging findings, showing a disrupted structural and functional maturation in VPT at TEA, to the regional gene expression profiles from mid- to late-gestation, we provide deeper insights into the potential biological mechanisms underlying preterm birth injury. Specifically, gliogenesis and pre-myelination processes may be particularly vulnerable to disruption following preterm birth, contributing to the observed regional alterations in cortical microstructure and functional BOLD variability.

## 4. Limitations

This study has limitations that should be considered. First, the sample size is modest. This limitation is largely related to the challenges associated with recruiting preterm infants and their families during a stressful time in their lives. The recruitment and execution of the project is further constrained by its longitudinal design, requiring two MRI scans, with an early first one by the 33rd wGA, followed by a second at TEA. This design necessitates the preterm infant to be clinically stable at 33 wGA, to undergo the MRI, which may not be always the case.

Second, the spatial resolution limitations of both functional and diffusion imaging may affect the accuracy of the registration of the RS-fMRI ICA-based brain atlas - originally in the subjects’ anatomical T2 space - to the functional or diffusion subjects’ spaces. Such may introduce partial volume effects. However, since the pipeline is automated, any potential systematic errors would be consistent across subjects, allowing for valid longitudinal comparisons and between-groups analysis.

Third, the clinical significance behind brain BOLD variability and cortical microstructural alterations identified by DKI, DTI and SMT metrics, following preterm birth, remains to be investigated. Evaluation of the long-term neurodevelopmental follow-up of this cohort during childhood is planned to investigate the relevance of these early imaging biomarkers.

Fourth, despite efforts to minimize motion during acquisition, neonatal MRI is inherently susceptible to motion artifacts, since no sedation was used; and despite application of motion correction analysis techniques, the quality of the data can still be affected by this limitation.

Fifth, the ICA-based brain atlas used in both our longitudinal and cross-sectional analyses was derived from a combination of VPT infants scanned at 33 wGA and at 40wGA, as well as FT, representing our cohort of infants. Despite the variability of the brain and, possibly, of the RSNs during this period, this approach was intentional, aiming to capture the variability of these RSNs during this period, which was considered essential for the longitudinal analysis. The same atlas was also applied to the cross-sectional analysis at term age, to ensure consistency in the regions evaluated, thereby facilitating direct comparisons with the longitudinal findings.

## 5. Conclusion

Our findings support the biological significance of brain’s BOLD variability during early preterm infants’ development, whose increase is accompanied by markers of regional cortical microstructural maturation and upregulation of genes mediating gliogenesis and pre-myelination events, specifically in primary sensory, sensorimotor and proto-DMN networks. Additionally, preterm birth affects these maturational processes, resulting in decreased cortical functional and microstructural maturity at term-equivalent age, compared to full-term birth.

By bridging genetic signatures with large-scale cortical organization, this work provides further insights regarding mechanisms underlying cortical maturation during late gestation and the putative mechanisms of preterm injury.

## 6. Materials and Methods

### 6.1. Subjects and study design

54 VPT infants (<32 wGA at birth) and 24 FT infants were recruited at the neonatal and maternity units of the University Hospitals of Geneva (HUG), Switzerland, from 2017 to 2020. Research Ethics Committee approval was granted and written parental consent was obtained prior to infant’s participation to the study.

Exclusion criteria for all babies included major brain lesions detected on the MRI, such as high-grade intraventricular hemorrhage, leukomalacia, as well as micro or macrocephaly or congenital syndromes.

For FT newborns, inclusion criteria included birth after 37 wGA, height, weight and head circumference above the 5th and below the 95th percentiles, APGAR score > 8 at 5 minutes and absence of resuscitation, infection or admission to the NICU (neonatal intensive care unit).

Six VPT infants were excluded from the study due to parental withdrawal or detection of a genetic anomaly, and two FT infants were excluded due to parental withdrawal.

Out study comprises both longitudinal and cross-sectional designs. The longitudinal design involves only our VPT infant cohort, who underwent MRI examinations at two time-points, firstly during the 33rd week of GA, and secondly at TEA. The cross-sectional design involves both the VPT at TEA and the FT newborns, who underwent a single MRI examination few days after being born (term age).

Infants whose MRI protocol acquisition was incomplete (not comprising a T2-weighted image, resting-state fMRI (RS-fMRI) sequence and multi-shell diffusion imaging (MSDI) sequence), without both longitudinal time-points (in preterm infants’ case) or whose images presented excessive motion were excluded from the analysis. The final sample comprised 46 VPT infants and 21 FT newborns, of which 31 VPT and 19 FT newborns had data suitable for the RS-fMRI analysis, while 39 VPT infants and 18 FT newborns had data suitable for the diffusion microstructural analysis, with an overlap of 24 VPT infants and 16 FT newborns between the two analyses. The flow chart of the participant selection process is provided in supplementary figure S5.

### 6.2. MRI acquisition

All infants were scanned after receiving breast or formula feeding during natural sleep (no sedation used). At 33 wGA, preterm infants were scanned using an MR compatible incubator (Lammers Medical Technology, Lübeck, Germany) and monitored using a Philips MR patient monitor Expression MR400 and Philips quadtrode MRI compatible neonatal ECG electrodes. At TEA, infants were scanned using a vacuum mattress for immobilization. All infants, at both time points, were monitored for heart rate and oxygen saturation during the entire scanning time. MR-compatible headphones (MR confon, Magdeburg, Germany) were used to protect the infants from scanner noise. MRI acquisition at both time-points was performed on a 3.0T Siemens Magnetom MR scanner (Siemens, Erlangen, Germany), using a 16-channel neonatal head coil. T2-weighted images were acquired using the following parameters: 113 coronal slices, TR= 4990 ms, TE= 160 ms, flip angle= 150°, matrix size= 256 × 164; voxel size= 0.8 × 0.8 × 1.2 mm^3^. RS-fMRI data acquisition was obtained by means of T2*-weighted gradient echo echo-planar imaging (EPI) sequence with the following parameters: 590 images, TR = 700 ms, TE = 30 ms, 36 slices, voxel size = 2.5 x 2.5 x 2.5 mm^3^, flip angle = 60°, multiband factor = 4. Multi-shell diffusion imaging (MSDI) was acquired with a single-shot spin echo echo-planar imaging (SE-EPI) Stejksal-Tanner sequence with the following parameters TE=85 ms, TR=3170 ms, voxel size 1.8 ×1.8 ×1.8 mm^3^, multi-band acceleration factor of 2, GRAPPA 2. Images were acquired in the axial plane, in anterior-posterior (AP) phase encoding (PE) direction, with 4 volumes without diffusion-weighting (b0); 10 non-collinear directions with b= 200 s/mm^2^, 30 non-collinear directions with b= 1000 s/mm^2^; 50 non-collinear directions with b= 2000 s/mm^2^. Additional b0 images were collected with reversed phase-encode blips, AP and posterior-anterior (PA), resulting in pairs of images with distortions going in opposite directions.

### 6.3. RS-fMRI analysis

#### 6.3.1. Pre-processing

The RS-fMRI data were preprocessed using SPM12 (Wellcome Department of Imaging Neuroscience, University College London, United Kingdom). The preprocessing steps included realignment to the mean functional volume, adjusting for motion, co-registration to the time-point specific structural image, alignment in Montreal Neurological Institute (MNI) space, and normalization, in which all the scans are warped to our cohort 40 wGA template. Slice-timing correction was omitted due to the fast TR of 700 ms used. Finally, the data were smoothed with a Gaussian filter of full width at half maximum (FWHM) of 5 mm. All volumes with a frame-wise displacement (Power, Mitra et al. 2014) greater than 0.5 mm or with a rate of BOLD signal changes across the entire brain (DVARS) greater than 3% were removed, along with the two previous and the two subsequent images. The remaining images were included for further analysis. A minimum of 50% of volumes remaining was set as a sufficient criteria for inclusion.

#### 6.3.2. Resting-state atlas creation

Resting-state networks (RSNs) were generated from our cohort of infants, comprising both VPT at 33- and 40 wGA, as well as the FT infants, and combined into a brain cortical parcellation atlas. To extract independent spatial networks, a group-level independent component analysis (group-ICA) was conducted; a z-score threshold of 2 was applied to these components. The results were acquired in a single group-level ICA using the GIFT toolbox in MATLAB (http://mialab.mrn.org/software/gift/index.html). Voxels in the cerebrospinal fluid, ventricles, eyes, and extracerebral areas (skull) were removed using a brain mask. The ICA was repeated 20 times using ICASSO for stability of the decomposition and determined 13 robust components when combining the 33- and 40 wGA scans, VPT and FT infants combined (supplementary figure S6). Among the 13 components, 3 reflected areas related to motion and blood vessels, rather than resting-state activity. These were considered noise and excluded from further analyses. The 10 remaining components were combined into a brain atlas. This group-level ICA-based atlas was manually corrected to ensure anatomical precision. The cerebellum and the brainstem were removed from the final atlas, in order to keep only the cortical and subcortical structures for further analysis, which resulted in 9 RSNs: Limbic (amygdala, insula, anterior cingulate cortex [ACC]), Paralimbic belt (orbitofrontal cortex [OFC], temporal pole), Thalamus, Prefrontal cortex [PFC], Visual, Auditory, Posterior cingulate cortex [PCC], Sensorimotor and Precuneus (Figure 6).

**Figure 6.**
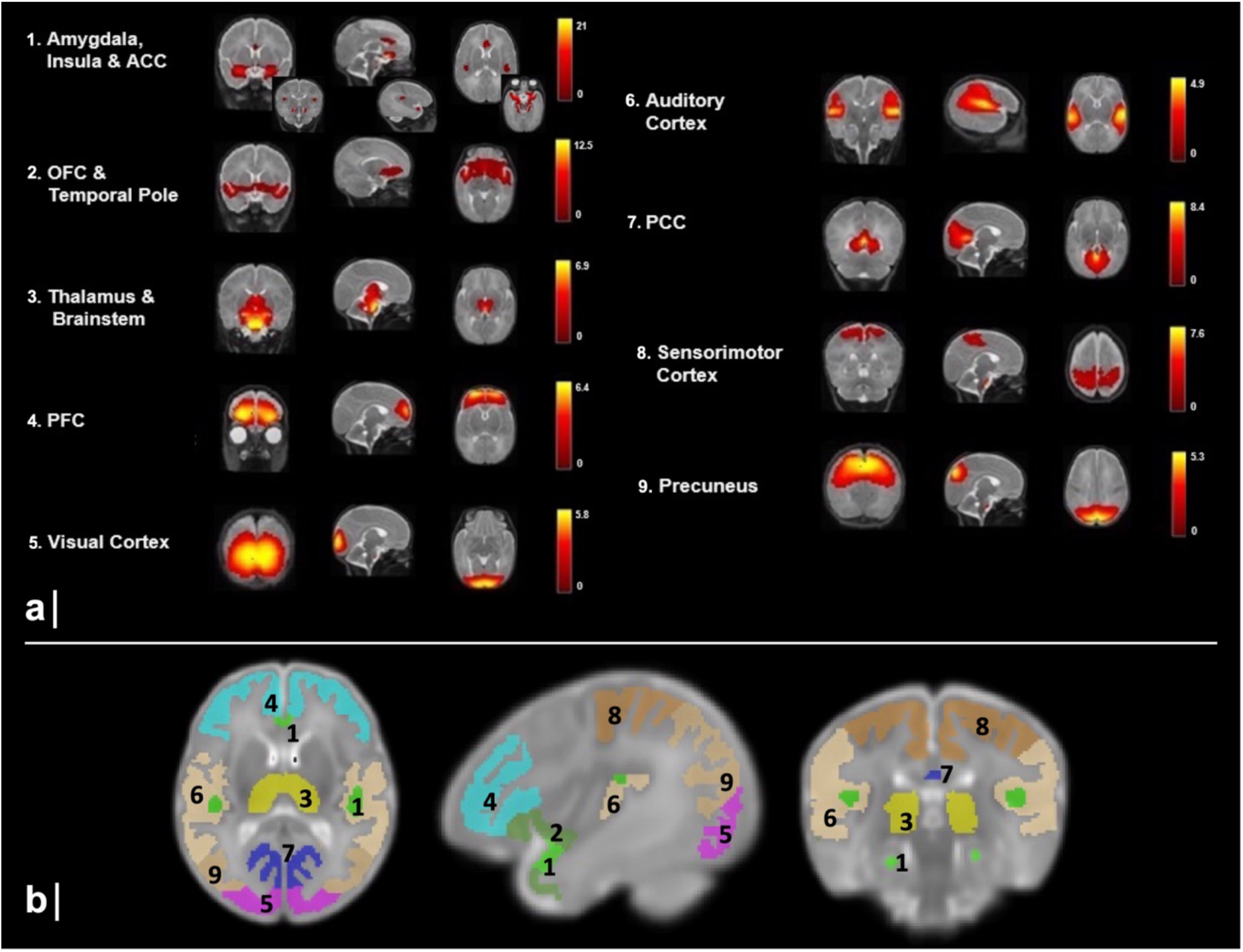
a) Main 9 components obtained from the data-driven ICA group analysis. Each row shows sagittal, coronal, and axial view of the components overlaid onto the 40 wGA template. A threshold at a z-score of 2 was applied. The colored bars show the corresponding z-score. **b) ICA-based brain atlas comprising 9 RSNs.** RSNs are illustrated in the T2w 40 wGA template, after GM masking. 1. Limbic (amygdala, insula, anterior cingulate cortex), 2. Paralimbic belt (orbitofrontal cortex, temporal pole), 3. Thalamus, 4. Prefrontal cortex, 5. Visual, 6. Auditory, 7. Posterior cingulate cortex, 8. Sensorimotor, 9. Precuneus.

#### 6.3.3. Regional cortical BOLD variability estimation

To extract regional time-series from the RS-fMRI data at each time-point, we used our ICA-based brain atlas consisting of 9 RSNs. First, the atlas was registered to each subject’s native time-point specific space (structural T2-weighted) using the diffeomorphic symmetric image normalization algorithm with cross-correlation as similarity metric (SyN-CC), from the Advanced Normalization Tools (ANTs) (Avants et al., 2011) toolbox.

Each subject’s structural T2-weighted image was segmented using the tissue segmentation maps from the UNC neonatal AAL atlas (Shi et al., 2011), which include Gray Matter (GM, cortical and deep gray matter), White Matter (WM) and Cerebral Spinal Fluid (CSF). These maps were first registered to the subject’s native space using the same ANTS algorithm described above, to obtain the three tissue-probability maps (TPMs) in the native space. Next, using these TPMs, the subject’s T2-weighted image was segmented using SPM12, so that subject-specific GM, WM and CSF probability maps were obtained in the subject’s T2 structural space.

To ensure that the BOLD signal was extracted only from GM voxels, the ICA-based brain atlas in the native space was GM-masked using the subject-specific TPMs obtained from the segmentation step. If the probability of one voxel being GM was less than the probability of this voxel being WM or CSF (P(GM) < P(WM) or P(GM) < P(CSF)), then this voxel was not considered as GM and was excluded from the atlas.

The subject’s GM-masked ICA-based atlas in the subject’s structural space was then registered to the subjects’ functional (BOLD) space and the average regional BOLD time-courses were extracted, after performing voxel-wise nuisance regression (CSF, WM, motion signals), detrending and smoothing (Gaussian FWHM 5mm), yielding a matrix with dimensions [#volumes, 9] for each subject and each RS-fMRI time-point (33 and 40 wGA). The 9 regional time-courses were bandpass filtered ([0.01Hz – 0.1Hz]) to discard noise components and non-resting state fMRI components. Finally, the BOLD signal variability (BOLD SD) was estimated by computing the sample standard deviation of the time-course of each of the 9 RSNs. This step resulted in 9 regional BOLD variability values per subject and per time-point.

### 6.4. Multi-Shell Diffusion Imaging analysis

#### 6.4.1. Pre-processing

MSDI data were preprocessed using MRtrix3 (version 3.0rc3, https://www.mrtrix.org) (Tournier et al., 2019) for denoising, bias field corrections and intensity normalization. The FSL diffusion toolbox (v5.0.11, https://fsl.fmrib.ox.ac.uk/fsl/fslwiki/) (Behrens et al., 2003, Smith et al., 2004) was used for preprocessing of MSDI data. FSL’s TOPUP (Andersson et al., 2003, Smith et al., 2004) was used to estimate the off-resonance field, which was then used as input for FSL’s EDDY function optimized for neonatal diffusion data, correcting for distortions induced by susceptibility and eddy currents, as well as by motion-induced signal dropout and intra-volume subject movement (Andersson et al., 2016, Andersson et al., 2017, Bastiani et al., 2019). Data were visually inspected to assure quality of motion artifacts correction.

#### 6.4.2. Diffusion cortical microstructural maps

After pre-processing, MSDI data was analyzed using three different methodological approaches.

To examine cortical microscopic features unconfounded by the directional structure, we use the Spherical Mean Technique (SMT) (Kaden et al., 2016), which focus on the direction-averaged diffusion weighted signal, independent of the fiber orientation distribution. MSDI was analyzed using the multi-compartment microscopic diffusion model from the publicly available SMT code (https://github.com/ekaden/smt). SMT metrics corresponding to intra-neurite volume fraction (intra), intrinsic diffusivity (Diff), extra-neurite mean diffusivity (extraMD) and extra-neurite transverse diffusivity (extraTrans) were calculated.

Alongside, we calculated Diffusion Tensor Imaging (DTI) metrics (fractional anisotropy, FA; mean diffusivity, MD) measuring Gaussian diffusion properties of the water in the tissue, as well as Diffusion Kurtosis Imaging (DKI) metrics (mean/radial/axial kurtosis, MK/RK/AK), evaluating Non-Gaussian diffusion and allowing the characterization of tissue microstructural heterogeneity (Jensen & Helpern, 2010; Jensen et al., 2005). For both DTI and DKI metrics, we used the code publicly available from DIPY library (https://docs.dipy.org/stable/interfaces/reconstruction_flow.html), applying the DKI model to the full MSDI data.

The GM-masked ICA-based brain atlas in the subject’s structural space was registered to the subjects’ dMRI space using ANTs (Avants et al., 2011), and averaged diffusion metrics were extracted per cortical RSN and per subject, at each time-point.

### 6.5. Gene expression analysis

Regional patterns of gene expression in the fetal cortex across gestation were obtained from the BrainSpan developmental transcriptomic dataset (http://development.psychencode.org/), comprising mRNA-seq collected from n = 41 specimens aged between 8 pcw and 40 postnatal years. For our study we have included data sampled from 18 post-mortem prenatal brain specimens, aged 8-37 pcw (n = 165 regional tissue samples, mean [SD] age = 17.74 [7.54] pcw, 53% male, mean [SD] postmortem interval = 6.52 [11.03] hours, mean [SD] RNA integrity number [RIN] = 9.34 [0.66]).

Tissue had been collected after obtaining parental or next of kin consent and with approval by the institutional review boards at the Yale University School of Medicine, the National Institutes of Health, and at each institution from which tissue specimens were obtained. For each brain, regional dissection including 11 neocortical regions was performed (dorsolateral PFC [DLPFC], ventrolateral PFC [VLPFC], OFC, medial frontal cortex [MFC], primary motor/sensory/auditory/visual cortex [M1, S1, A1C, V1], inferior parietal cortex [IPC], superior temporal cortex [STC] and inferior temporal cortex [ITC]). Tissue processing and detailed anatomical boundaries for each cortical region at each stage of development are provided elsewhere (Kang et al., 2011; Li et al., 2018). Regional tissue samples were subject to mRNA-seq using an Illumina Genome Analyzer IIx (Illumina, San Diego, California, United States of America) and mRNA-seq data processed using RSEQtools (version 0.5) (Habegger et al., 2011). Gene expression was measured as RPKM. Conditional quantile normalization was performed to remove guanine-cytosine (GC) content bias, and ComBat was used to remove technical variance due to processing site (Yale or University of Southern California) (Hansen et al., 2012; Johnson et al., 2007; Li et al., 2018).

The prenatal gene expression data were initially filtered to only include protein-coding genes (NCBI GRCh38.p12, n = 18,524 out of a possible 20,720). In order to restrict our analysis to focus on genes expressed in the developing cortex, this list was further filtered to only contain genes expressed by cells in the fetal cortex based on the composite list of prenatal cell markers from 5 independent single-cell RNA studies of the developing fetal cortex (Ball et al., 2020). This resulted in expression data from a final set of 5,287 genes.

For our study, gestational periods were clustered into early-fetal (T0, 8-13 pcw), mid-fetal (T1, 16-22 pcw) and late-fetal (T2, 35-37 pcw). The neocortical regions with available genetic data were matched to our RSNs as follows: MFC = Limbic (1), OFC = Paralimbic belt (2), DLPFC+VLPFC = PFC (4), V1 = Visual (5), A1C = Auditory (6), M1+S1 = Sensorimotor (8), IPC = Precuneus (9). Next, these regions were clustered according to the presence (cluster 1) or absence (cluster 2) of significant BOLD variability changes from 33- to 40-wGA (Figure 4a-b).

### 6.6. Statistical analysis

#### 6.6.1. Neonatal and demographic data

To test for differences in demographic and perinatal data between VPT and FT groups, categorical variables (sex, intrauterine growth restriction, sepsis, intraventricular hemorrhage grade I) were analyzed using chi-squared test, whereas continuous variables were compared using independent samples t-test, with group as an independent variable and the following dependent variables: GA at birth, GA at MRI at TEA, birth weight, birth height, birth head circumference, APGAR score at 1 and 5 min after birth and parental socio-economic status score (SES) (Largo et al., 1989).

#### 6.6.2. MRI data analysis

For each preterm participant and per time point (33- and 40-wGA), regional imaging metrics (BOLD variability and microstructural diffusivity) were scaled to the maximum regional absolute change, preserving the original directionality of changes. The metrics and averaged across participants to produce a group average region x metric matrix representing the relative variation of each imaging metric across the cortical RSN regions.

Longitudinal (from 33- to 40-wGA) and cross-sectional differences (at TEA, between VPT at 40-wGA and FT newborns) in both cortical BOLD variability and microstructural diffusivities were assessed with paired-samples t-test and independent samples t-test, respectively, with FDR correction for multiple comparisons.

Microstructural diffusivities were clustered using the R-package and function ConsensusClusterPlus (k-means base algorithm and Euclidean distance) (Wilkerson & Hayes, 2010), to identify stable clusters of regional diffusivities. The optimal number of clusters was determined using the elbow method, based on the within-cluster sum of squares (WCSS). This approach enabled grouping cortical regions based on the similarity of their longitudinal microstructural maturation, and to compare these microstructural clusters with those derived from BOLD variability longitudinal changes.

Lasso regression was used to identify the most relevant microstructural predictors of BOLD variability longitudinal changes.

#### 6.6.3. Gene expression data analysis

A linear mixed effects regression model (LMM) was used to identify significant changes in gene expression as a function of regional cluster (1 and 2) and time, with fixed effects of sex and RIN (RNA integrity number), including specimen ID as a random effect. We identified genes with a significant cluster*time interaction and calculated estimated marginal means, followed by pairwise comparisons to assess differences in gene expression over time and between clusters. Multiple comparisons were adjusted using false discovery rate (FDR) correction. Modelling was performed in R (4.4.1) using “lmer()” function from lme4 package. Gene function was classified using gene set enrichment analysis (GSEA) using WebGestalt (Wang et al., 2017). Enrichment ratios were calculated as the proportion of cell class-specific genes in our gene list of interest compared to the proportion in the full background set. The background gene set was defined as the full list of protein-coding genes included in the analysis (n = 5,287 expressed by cells in the fetal cortex). We corrected for multiple comparisons across cell classes using FDR. The genes with a significant cluster*time interaction, found to have a significant enrichment ratio linked to neocortical organization, were combined for expression analysis across time points (T0, T1 and T2) and between clusters (1 and 2) using two-way ANOVA followed by post-hoc comparisons using Tukey’s test (HSD).

## Data and code availability

All neuroimaging data were acquired in the context of a research project approved by the ethical committee in 2016. The patient consent form did not include any clause for reuse or sharing of data. Software and code used in this study are publicly available as part of FSL v5.0.10 (https://fsl.fmrib.ox.ac.uk/fsl/fslwiki/), MRtrix3 (Tournier et al., 2019), SMT (https://github.com/ekaden/smt) and DIPY (https://docs.dipy.org/stable/interfaces/reconstruction_flow.html) software packages. dMRI data were pre-processed using EDDY command adapted for neonatal motion, from the neonatal dMRI automated pipeline from developing Human Connectome Project (dHCP, http://www.developingconnectome.org), and can be found at: https://git.fmrib.ox.ac.uk/matteob/dHCP_neo_dMRI_pipeline_release (Bastiani et al., 2019). Developmental RNA-seq data were accessed via: http://development.psychencode.org/.

## Acknowledgements

The authors thank all clinical staff, namely in the neonatology and in the unit of development of the HUG Children’s Hospital, all parents and newborns participating in the project, the Pediatrics Clinic Research Platform and the Center for Biomedical Imaging (CIBM) of the University Hospitals of Geneva, for all their valuable help and support.

## Author contributions

**J.S.A.:** data collection and investigation, conceptualization and design of methods and analysis, formal analysis, writing - original draft. **A.B.:** formal analysis, design of methods and analysis, writing - review & editing. **S.L:** conceptualization and design of methods and analysis, writing – review & editing. **E.F.:** conceptualization of methods, writing - review & editing. **A.V.:** data analysis, writing - review & editing. **L.L**: data analysis, writing - review & editing. **S.C.:** resources, investigation. **F.L.:** resources, investigation. **D.V.V.:** conceptualization and design of methods and analysis; writing - review & editing. **G.B.:** conceptualization and design of methods and analysis, supervision, resources, writing - review & editing. **P.S.H.:** conceptualization, supervision, resources, funding acquisition, writing - review & editing.

