## Supplementary material for "Regional BOLD variability reflects microstructural maturation and neuronal ensheathment in the preterm infant cortex": Figure S1

**Figure S1. Longitudinal changes in premature infants' microstructural diffusivities, from 33- to 40-wGA, averaged per RSN and scaled by the largest observed change per metric.** a-b) Microstructural DKI metrics delta (Δ) change, including mean (MK), radial (RK) and axial kurtosis (AK); c-d) DTI metrics delta (Δ) change, including mean diffusivity (MD) and fractional anisotropy (FA); e-f) SMT metrics delta (Δ) change, including intra-neurite volume fraction (intra), intrinsic diffusivity (diff), extra-neurite mean diffusivity (extraMD) and extra-neurite transverse diffusivity (extraTrans).

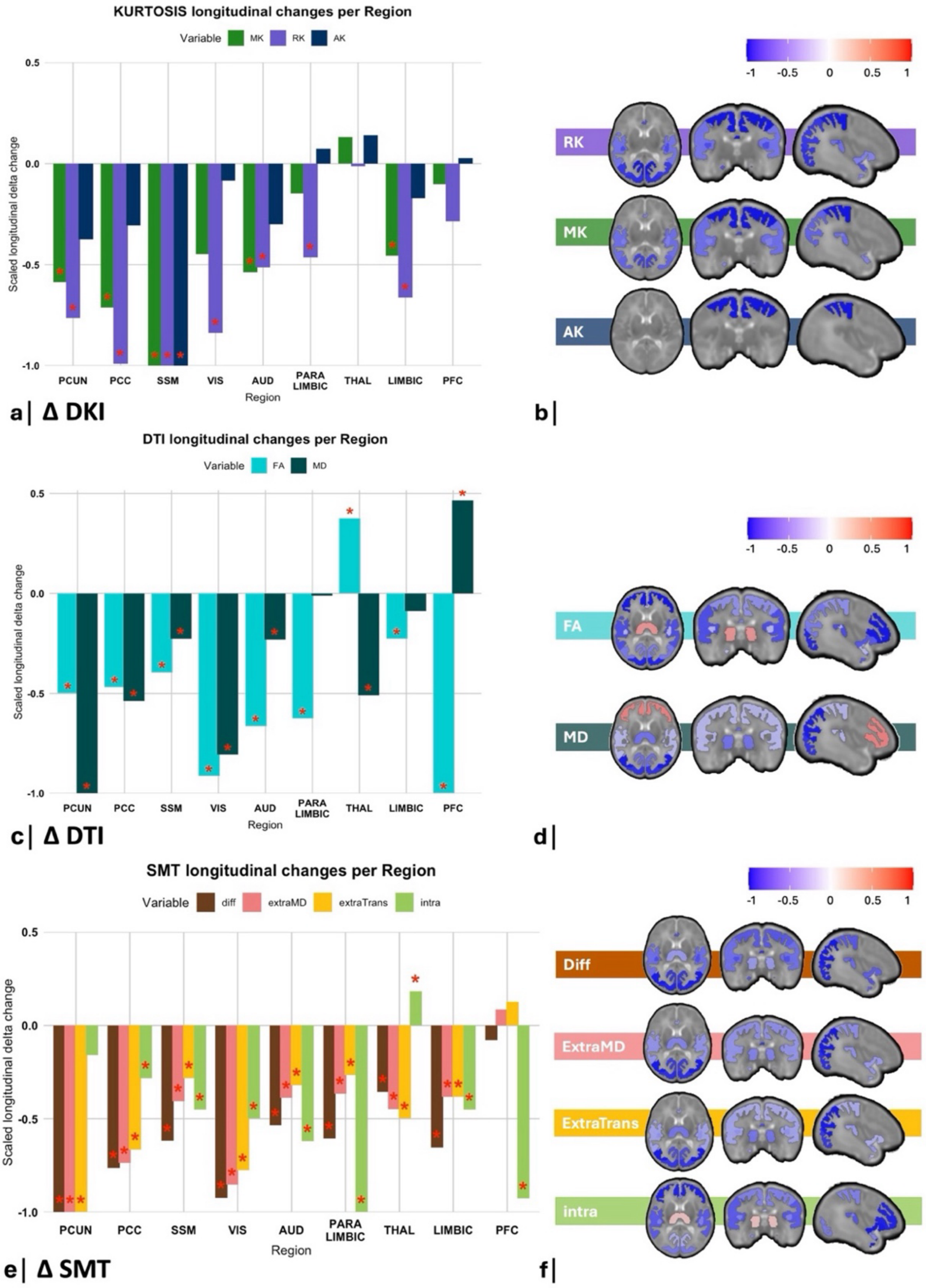
