## Supplementary material for "Regional BOLD variability reflects microstructural maturation and neuronal ensheathment in the preterm infant cortex": Figure S2

**Figure S2. Results from the consensus clustering of the combined 9 distinct diffusion microstructural measures delta changes over time.** a) Heatmap depicting the regional maturational patterns from 33- to 40-wGA, with rows and columns representing the RSNs and color intensity in each cell reflecting the composite changes over time of the microstructural metrics. b) Composition of the identified 3 regional patterns of microstructural delta changes (pattern 1: SSM; pattern 2: Visual, PCC and PCUN; pattern 3: PFC, Paralimbic, Limbic, Thalamus and Auditory). c) Brain plots illustrating the 3 distinct regional patterns depicted by the consensus clustering analysis, color-coded according to regional pattern (red = pattern 1, orange = pattern 2, blue = pattern 3).

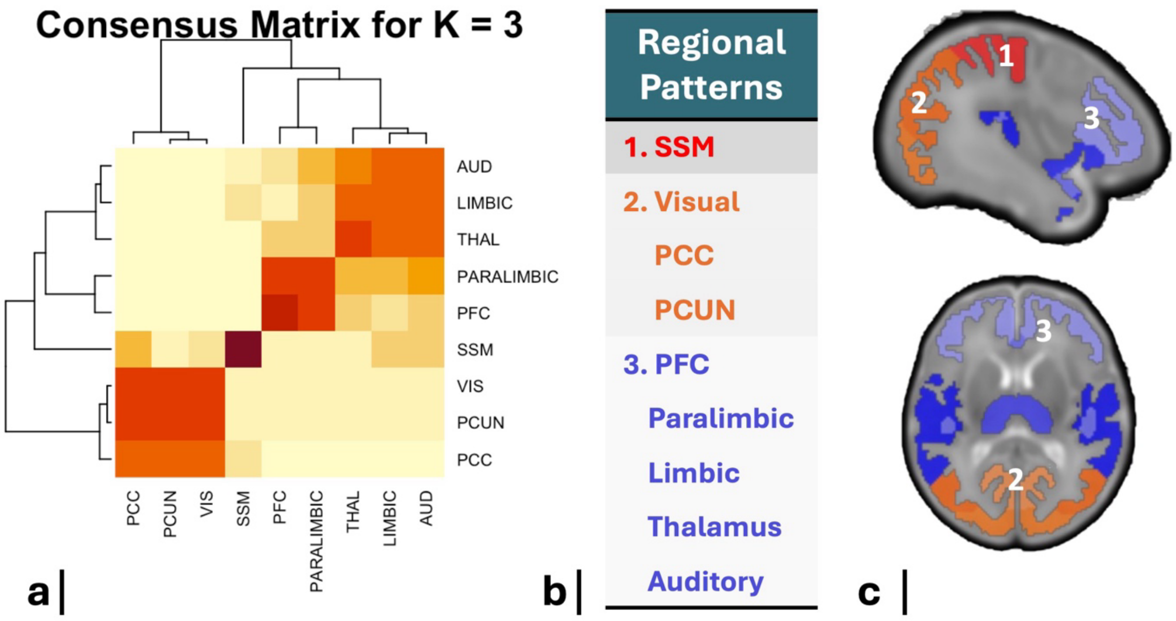


To better understand the spatio-temporal patterns of cortical microstructural maturation across the various RSNs, occurring from 33- to 40-wGA, we have combined the longitudinal changes of all the diffusion microstructural measures across the different dMRI models (DTI, DKI and SMT) and performed a consensus clustering analysis (k=3).

Microstructural diffusivities were clustered using ConsensusClusterPlus function (k-means base algorithm and Euclidean distance) in R (4.4.1) , to identify stable clusters of regional diffusivities. The optimal number of clusters was determined using the elbow method, based on the within-cluster sum of squares (WCSS).

Three distinct regional microstructural maturational patterns were identified, aligning with the central-to-peripheral and posterior-to-anterior known gradients in brain maturation, aligning with the brain myelination order of Kinney (Kinney et al., 1988). Microstructure cluster 1 and 2 are similar to BOLD SD cluster 1, whereas Microstructure cluster 3 is similar to BOLD SD cluster 2.
