## Supplementary material for "Regional BOLD variability reflects microstructural maturation and neuronal ensheathment in the preterm infant cortex": Table S3

**Table S3. Gene Ontology enrichment of genes that increased significantly more in cluster 1 than 2 from mid- to late-fetal period**, using background reference set (5,287 genes).

| | | Reference set | | |
|---|---|---|---|---|
| | | Fetal gene markers (n=5287) | | |
| GO term[†] | Description | Enrichment | FDR | Genes |
| GO:0007272 | ensheathment of neurons | 6.15 | 0.00001* | CD9; CLDN11; COL6A1; ERBB3; MAG; MAL; MBP; MOBP; NKX6-2; PLLP; PLP1; PMP22; TNFRSF1B; UGT8 |
| GO:0042063 | gliogenesis | 2.85 | 0.01796* | APCDD1; CD9; COL6A1; ERBB3; GPR17; GPR183; GPR37L1; IL6ST; MAG; MAL; MOBP; NKX6-2; PLP1; TNFRSF1B |
| GO:0043062 | extracellular structure organization | 3.58 | 0.01796* | CAV1; CAV2; COL18A1; COL5A2; COL6A1; EFEMP2; LAMC1; SLC39A8; TNFRSF1B; VWA1 |
| GO:0045229 | external encapsulating structure organization | 3.58 | 0.01796* | CAV1; CAV2; COL18A1; COL5A2; COL6A1; EFEMP2; LAMC1; SLC39A8; TNFRSF1B; VWA1 |
| GO:0019221 | cytokine-mediated signaling pathway | 3.61 | 0.00797* | CAV1; CD74; F3; GPR17; IFI27; IL6ST; IRF1; MX1; OAS2; PADI2; PTPRC; SP100; TNFRSF1B |
| GO:0050817 | coagulation | 4.10 | 0.01351* | ACTN1; CAV1; CD9; CSRP1; F3; IL6ST; MMRN1; NFE2L2; PAPSS2; STXBP3 |
| GO:0042060 | wound healing | 3.04 | 0.01365* | ACTN1; CAV1; CD9; CSRP1; ELK3; ERBB3; F3; GATA2; HBEGF; IL6ST; MMRN1; NFE2L2; PAPSS2; STXBP3 |
| GO:0097191 | extrinsic apoptotic signaling pathway | 3.84 | 0.01548* | BAG3; CAV1; ERBB3; IFI27; MAL; PTPRC; SH3RF1; SP100; STK3; TNFRSF1B |
| GO:0033002 | muscle cell proliferation | 3.86 | 0.01796* | APOD; CALCRL; CAV2; EFEMP2; HBEGF; IGFBP5; JUN; NDRG2; PTGS2 |
| GO:0050878 | regulation of body fluid levels | 3.55 | 0.00797* | ACTN1; CAV1; CD9; CSRP1; EMP2; F3; IL6ST; MMRN1; NFE2L2; OAS2; PAPSS2; STXBP3; WFS1 |

*p<0.05 after correction for multiple comparisons

[†] top 10 terms with ‘fetal gene markers’ background set are listed
