## Supplementary material for "Regional BOLD variability reflects microstructural maturation and neuronal ensheathment in the preterm infant cortex": Table S4

**Table S4. Effect of preterm birth on BOLD variability and cortical microstructure at TEA per region.** Metrics include: BOLD variability (BOLD SD), Mean Kurtosis (MK), Radial Kurtosis (RK), Axial Kurtosis (AK), Mean Diffusivity (MD), Extra-neurite Mean Diffusivity (ExtraMD), Extra-neurite Transverse Diffusivity (ExtraTrans), intrinsic diffusivity (Diff). Numbers in bold indicate significant results between groups that survived FDR correction ($p \leq 0.05$).

| **Cortical region** | **Metric** | **Mean measurement (95% CI)** | | **Group-effect** |
|---|---|---|---|---|
| | | **VPT** | **FT** | **p-value (FDR adjusted)** |
| **Precuneus** | BOLD SD | 16.5 (14.3-18.8) | 17.6 (14.9-20.3) | 0.51 |
| | MK | 0.42 (0.39-0.45) | 0.37 (0.33-0.41) | **0.039*** |
| | RK | 0.41 (0.37-0.43) | 0.33 (0.27-0.38) | **0.016*** |
| | AK | 0.46 (0.44-0.48) | 0.45 (0.42-0.47) | 0.66 |
| | MD | 0.0017 (0.0016-0.0018) | 0.0014 (0.0013-0.0015) | < **0.001***** |
| | ExtraMD | 0.0018 (0.0017-0.0019) | 0.0015 (0.0014-0.0016) | < **0.001***** |
| | ExtraTrans | 0.0017 (0.0016-0.0018) | 0.0014 (0.0013-0.0015) | < **0.001***** |
| | Diff | 0.0021 (0.0019-0.0022) | 0.0017 (0.0016-0.0018) | < **0.001***** |
| **Posterior Cingulate Cortex** | BOLD SD | 13.6 (12.1-15.0) | 18.6 (15.1-22.1) | **0.02*** |
| | MK | 0.43 (0.41-0.46) | 0.39 (0.36-0.42) | **0.03*** |
| | RK | 0.40 (0.37-0.43) | 0.33 (0.28-0.38) | **0.016*** |
| | AK | 0.50 (0.44-0.47) | 0.49 (0.47-0.51) | 0.66 |
| | MD | 0.0014 (0.0013-0.0015) | 0.0013 (0.0012-0.0014) | < **0.001***** |
| | ExtraMD | 0.00159 (0.0015-0.0016) | 0.0014 (0.0013-0.0015) | < **0.001***** |
| | ExtraTrans | 0.00149 (0.0014-0.0015) | 0.0013 (0.0012-0.0014) | < **0.001***** |
| | Diff | 0.0018 (0.0017-0.0019) | 0.0016 (0.0015-0.0017) | < **0.001***** |
| **Sensorimotor** | BOLD SD | 10.0 (8.6-11.5) | 12.9 (10.3-15.6) | 0.09 |
| | MK | 0.47 (0.44-0.50) | 0.42 (0.37-0.47) | **0.05*** |
| | RK | 0.44 (0.41-0.48) | 0.35 (0.29-0.41) | **0.015*** |
| | AK | 0.52 (0.50-0.55) | 0.52 (0.49-0.56) | 0.85 |
| | MD | 0.0018 (0.0017-0.0019) | 0.0013 (0.0012-0.0014) | < **0.001***** |
| | ExtraMD | 0.0019 (0.0018-0.002) | 0.00146 (0.0014-0.0015) | < **0.001***** |
| | ExtraTrans | 0.0018 (0.0017-0.0019) | 0.00136 (0.0013-0.0014) | < **0.001***** |
| | Diff | 0.0022 (0.0020-0.0023) | 0.0017 (0.0015-0.0018) | < **0.001***** |
| Visual | BOLD SD | 16.7 (14.9-18.5) | 23.1 (18.9-27.2) | **0.02*** |
| | MK | 0.42 (0.39-0.45) | 0.36 (0.32-0.40) | **0.03*** |
| | RK | 0.39 (0.35-0.42) | 0.31 (0.25-0.36) | **0.015*** |
| | AK | 0.48 (0.45-0.50) | 0.47 (0.45-0.48) | 0.66 |
| | MD | 0.0015 (0.0014-0.0015) | 0.0013 (0.0012-0.0014) | **0.005**** |
| | ExtraMD | 0.0016 (0.0015-0.0017) | 0.0015 (0.0014-0.0016) | **0.005**** |
| | ExtraTrans | 0.0015 (0.0014-0.0016) | 0.0014 (0.0013-0.0015) | **0.001**** |
| | Diff | 0.0018 (0.0017-0.0019) | 0.0017 (0.0015-0.0018) | **0.02*** |
| Auditory | BOLD SD | 10.7 (9.2-12.1) | 14.8 (12.3-17.4) | **0.02*** |
| | MK | 0.46 (0.44-0.49) | 0.41 (0.36-0.45) | **0.03*** |
| | RK | 0.45 (0.42-0.48) | 0.38 (0.30-0.42) | **0.015*** |
| | AK | 0.50 (0.48-0.52) | 0.48 (0.46-0.51) | 0.57 |
| | MD | 0.0016 (0.0015-0.0017) | 0.0014 (0.0013-0.0015) | < **0.001***** |
| | ExtraMD | 0.0018 (0.0017-0.0019) | 0.0015 (0.0014-0.0016) | < **0.001***** |
| | ExtraTrans | 0.00167 (0.0016-0.0017) | 0.00144 (0.0014-0.0015) | < **0.001***** |
| | Diff | 0.0021 (0.0020-0.0022) | 0.0017 (0.0016-0.0018) | < **0.001***** |
| Paralimbic | BOLD SD | 11.4 (10.1-12.8) | 13.2 (10.4-15.8) | 0.30 |
| | MK | 0.46 (0.43-0.48) | 0.39 (0.34-0.44) | **0.03*** |
| | RK | 0.43 (0.40-0.46) | 0.34 (0.28-0.40) | **0.015*** |
| | AK | 0.49 (0.47-0.52) | 0.47 (0.43-0.50) | 0.49 |
| | MD | 0.0017 (0.0016-0.0018) | 0.0014 (0.0013-0.0015) | < **0.001***** |
| | ExtraMD | 0.00185(0.0018-0.0019) | 0.00157 (0.0015-0.0016) | < **0.001***** |

| | | | | |
|---|---|---|---|---|
| | ExtraTrans | 0.0017 (0.0016-0.0018) | 0.00148 (0.0014-0.0015) | < **0.001*** ** |
| | Diff | 0.0021 (0.0020-0.0022) | 0.0018 (0.0017-0.0019) | < **0.001*** ** |
| Thalamus | BOLD SD | 11.5 (9.8-13.1) | 19.5 (16.5-22.5) | < **0.001*** ** |
| | MK | 0.38 (0.35-0.41) | 0.33 (0.28-0.38) | 0.099 |
| | RK | 0.33 (0.29-0.37) | 0.26 (0.20-0.33) | 0.06 |
| | AK | 0.44 (0.42-0.47) | 0.43 (0.39-0.47) | 0.66 |
| | MD | 0.00123 (0.0012-0.0013) | 0.00117 (0.0011-0.0012) | **0.007** |
| | ExtraMD | 0.00135 (0.0013-0.0014) | 0.00123 (0.0011-0.0012) | < **0.001*** ** |
| | ExtraTrans | 0.00127 (0.0012-0.0013) | 0.00117 (0.0011-0.0012) | < **0.001*** ** |
| | Diff | 0.0015 (0.0014-0.0016) | 0.0013 (0.0012-0.0014) | < **0.001*** ** |
| Limbic | BOLD SD | 10.7 (9.2-12.2) | 13.5 (10.8-16.3) | 0.11 |
| | MK | 0.44 (0.41-0.47) | 0.39 (0.34-0.43) | **0.03*** |
| | RK | 0.40 (0.37-0.43) | 0.32 (0.26-0.37) | **0.015*** |
| | AK | 0.51 (0.49-0.53) | 0.48 (0.46-0.51) | 0.49 |
| | MD | 0.00156 (0.0014-0.0016) | 0.00136 (0.0013-0.0014) | < **0.001*** ** |
| | ExtraMD | 0.00176 (0.0017-0.0018) | 0.0015 (0.0014-0.0016) | < **0.001*** ** |
| | ExtraTrans | 0.0016 (0.0015-0.0017) | 0.0014 (0.0013-0.0015) | < **0.001*** ** |
| | Diff | 0.0020 (0.0019-0.0021) | 0.0018 (0.0017-0.0019) | < **0.001*** ** |
| Prefrontal Cortex | BOLD SD | 16.8 (13.2-20.4) | 19.8 (15.6-24.0) | 0.30 |
| | MK | 0.47 (0.44-0.50) | 0.41 (0.36-0.46) | **0.03*** |
| | RK | 0.45 (0.41-0.48) | 0.35 (0.28-0.41) | **0.015*** |
| | AK | 0.51 (0.49-0.54) | 0.50 (0.47-0.54) | 0.66 |
| | MD | 0.00188 (0.0018-0.0019) | 0.0014 (0.0013-0.0015) | < **0.001*** ** |
| | ExtraMD | 0.0020 (0.0019-0.0021) | 0.00156 (0.0015-0.0016) | < **0.001*** ** |
| | ExtraTrans | 0.0018 (0.0017-0.0019) | 0.0014 (0.0013-0.0015) | < **0.001*** ** |
| | Diff | 0.0022 (0.0022-0.0024) | 0.00146 (0.0014-0.0015) | < **0.001*** ** |
