## Supplementary figures and images for "Regional BOLD variability reflects microstructural maturation and neuronal ensheathment in the preterm infant cortex"

### Figure S5

**Figure S5.** Flow chart of participant selection

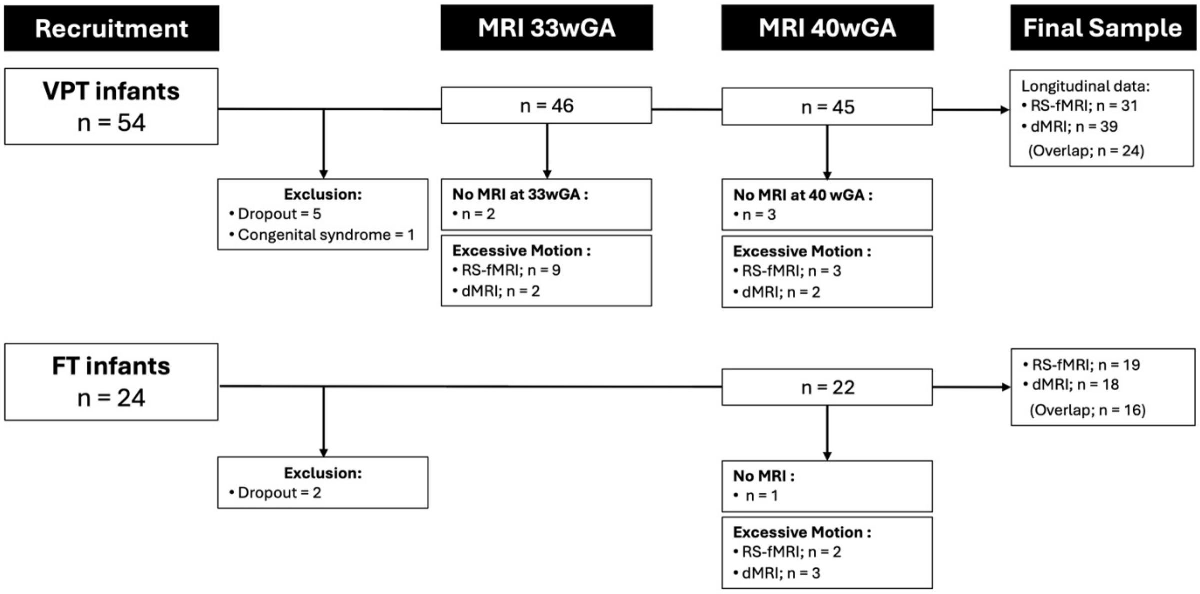

### Figure S6

**Figure S6.** Components obtained from the data-driven ICA group analysis

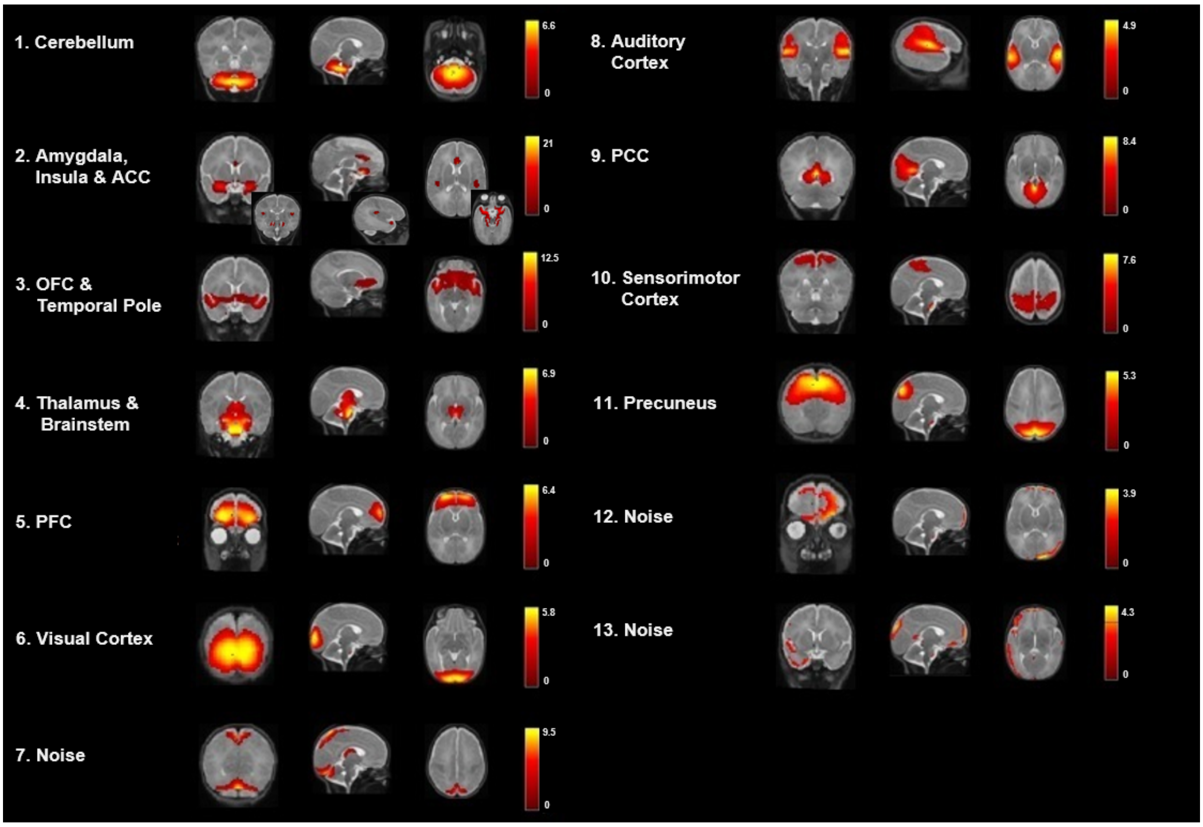
